# Hunting for extremophiles: a systematic screening of freshwater microalgae for tolerance to high pH and high alkalinity cultivation

**DOI:** 10.1101/2025.09.15.676395

**Authors:** Patrick K. Thomas, Robin Gerlach, Anita Narwani

**Author notes:** Current address of corresponding author: Patrick Thomas, Center for Biofilm Engineering, 366 Barnard Hall, Montana State University, Bozeman, MT, 59717, USA.

## Abstract

Microalgae hold the potential to supply sustainable food, fuel, plastics, and chemicals at commercial scales. Cultivating microalgae at extreme pH (>10) and high alkalinity provides multiple benefits including 1) reducing the risk of contamination by undesired organisms and 2) enabling direct air capture of CO_2_, which expands the land area suitable for algae farming compared to using CO_2_ point sources alone. However, we currently have a limited understanding of which algal taxa can grow under these conditions. Therefore, we conducted a high-throughput screening of 49 freshwater microalgae strains, comprising 40 species, for their ability to grow in moderate (pH 8.5, 25 mM alkalinity), high (pH 10, 75 mM alkalinity), and extreme (pH 10, 150 mM alkalinity) cultivation environments. Our results show that moderate alkalinity tends to significantly increase algae growth (including potentially harmful strains). However, higher levels inhibit all but a small subset of green algae and cyanobacteria. Effects of salinity and alkalinity differed, indicating they are broadly decoupled. Our results identify new industrially relevant alkaline-tolerant strains, show that algae isolated from “normal” ecosystems can be extremophilic, and suggest that future bioprospecting efforts for alkaline-tolerant algae adapted to local climatic conditions could yield additional productivity gains for the algae industry.

**Synopsis:** We tested if 49 distinct microalgae strains could grow in high pH environments and found new and productive alkaline-tolerant strains

**Abstract/TOC Graphic:** 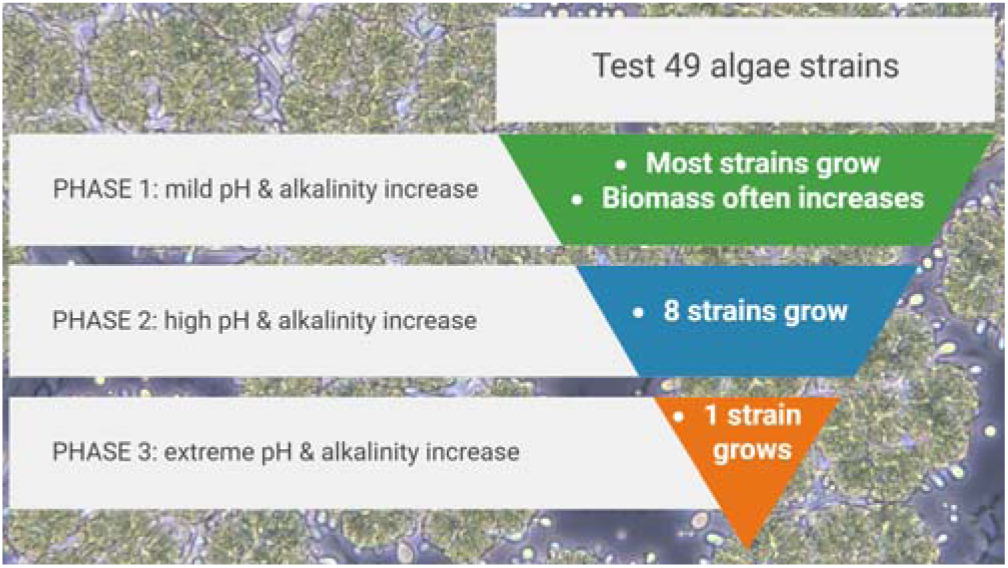

## Introduction

Microalgae farming has the potential to address several Sustainable Development Goals by provisioning much-needed protein and bio-based fuels, chemicals, and polymers to society while also stimulating rural economies and more efficiently using water resources^1^. In addition to the many foreseen benefits of developing commercial scale algae farms for food, feed, fuel, and bioproducts, there are also several challenges that have limited the economic viability of scaling up algae cultivation^2,3^. These challenges often involve overlapping limitations related to algal ecology and physiology, which can lead to yield instability, as well as process engineering bottlenecks, which in turn can hinder economic benefits of algae production. For example, outdoor algae cultures are subject to attack by “natural enemies” of algae including parasites, pathogens, and grazers, as well as invasion “weedy” algae strains^4–6^. Engineering barriers include the need to increase harvesting and conversion efficiency^7^, to sustainably source nutrients for algal fertilizers^8^, and to optimize the delivery of inorganic carbon to algae in a scalable and economical manner^9^. Proposed solutions for some of these challenges include: pest management via the use of extreme pH/salinity/temperatures^10^, the addition of biocides^11^, manipulation of algal microbiomes^12,13^, or the use of diverse algal polycultures that are reported to be less prone to pest outbreaks^14,15^.

Of these proposed solutions, cultivation at high pH and high alkalinity in particular holds promise, largely because it appears to have multiple co-benefits that address both biological and process engineering constraints for algae farming^16,17^. Specifically, high pH cultivation (i.e., > pH 10) excludes many potential biological pests from the system, as most grazers and other harmful species are not adapted to such extreme pH conditions. At the same time, using high alkalinity ponds allows algae farming to be decoupled from point sources of CO_2_; this is because the high availability of inorganic carbon in high carbonate alkalinity solutions combined with high mass transfer rates of CO_2_ from the atmosphere facilitates direct air capture of carbon. Examples of demonstrated high productivity in commercially-relevant strains under high pH/alkalinity include the green algae *Chlorella* sp. SLA-04^18,19^ and *Dunaliella salina*^20^, the diatom *Nitzschia inconspicua* str. Hildebrandi^21^, and several cyanobacteria ranging from well-known commercial strains like *Spirulina*^22^ to model strains like *Synechocystis* PCC 6803^20^ and a newly isolated strain of *Cyanobacterium* from an alkaline lake^23^.

Despite the potential of high pH and high alkalinity cultivation to enhance the reliability and productivity of sustainable algae farming, only a relatively limited number of algae strains have been tested for their ability to grow in alkaline conditions, leaving us with a minimal understanding of the algal taxa that may tolerate such cultivation environments. Moreover, while researchers have identified several economically beneficial alkaliphilic strains, we currently lack an understanding of whether potentially harmful algae strains could also grow in high pH and high alkalinity ponds. In this context, “harmful” refers to any microalgae species which would cause economic harm to algae farms (e.g., those marked with an asterisk in Table 1 below). For example, toxin-producing cyanobacteria such as *Microcystis* and *Planktothrix*, which form harmful blooms in natural aquatic systems^24^, could threaten the safety of algae-based food crops, while harmful mixotrophic algae such as *Poterioochromonas* can directly consume smaller algal taxa, thus decimating crops^25^.

**Table 1.**
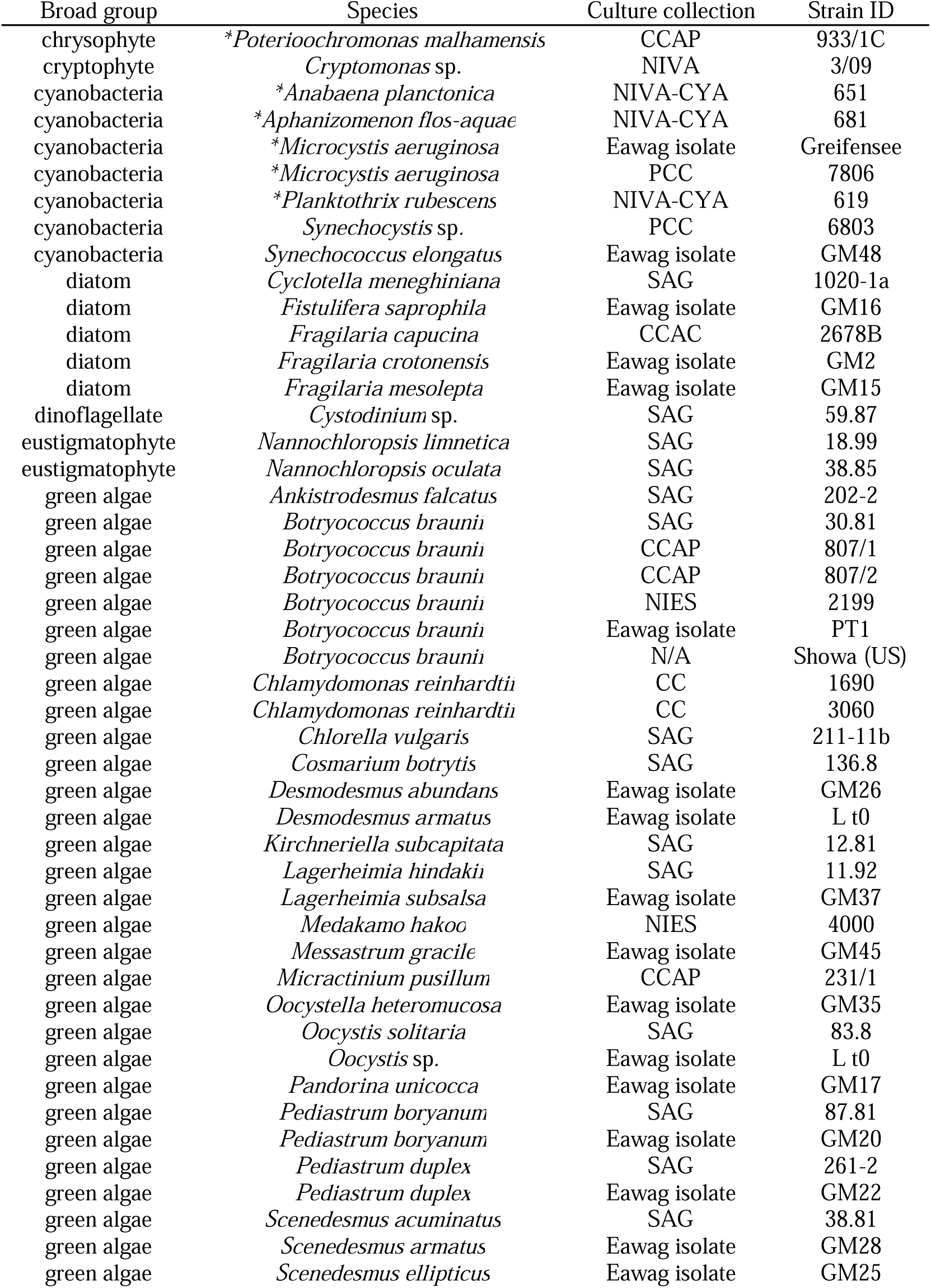
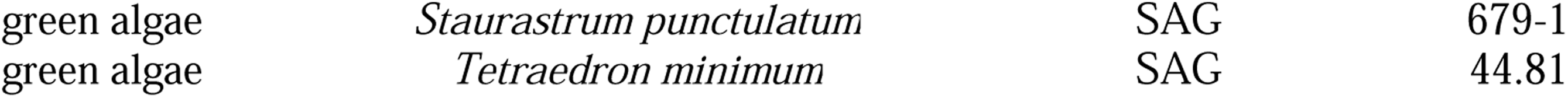
Algae strains used in this study. *Denotes strains designated as potentially harmful for commercial algae farming.

| Broad group | Species | Culture collection | Strain ID |
| --- | --- | --- | --- |
| chrysophyte | * *Poterioochromonas malhamensis* | CCAP | 933/1C |
| cryptophyte | *Cryptomonas* sp. | NIVA | 3/09 |
| cyanobacteria | * *Anabaena planctonica* | NIVA-CYA | 651 |
| cyanobacteria | * *Aphanizomenon flos-aquae* | NIVA-CYA | 681 |
| cyanobacteria | * *Microcystis aeruginosa* | Eawag isolate | Greifensee |
| cyanobacteria | * *Microcystis aeruginosa* | PCC | 7806 |
| cyanobacteria | * *Planktothrix rubescens* | NIVA-CYA | 619 |
| cyanobacteria | *Synechocystis* sp. | PCC | 6803 |
| cyanobacteria | *Synechococcus elongatus* | Eawag isolate | GM48 |
| diatom | *Cyclotella meneghiniana* | SAG | 1020-1a |
| diatom | *Fistulifera saprophila* | Eawag isolate | GM16 |
| diatom | *Fragilaria capucina* | CCAC | 2678B |
| diatom | *Fragilaria crotonensis* | Eawag isolate | GM2 |
| diatom | *Fragilaria mesolepta* | Eawag isolate | GM15 |
| dinoflagellate | *Cystodinium* sp. | SAG | 59.87 |
| eustigmatophyte | *Nannochloropsis limnetica* | SAG | 18.99 |
| eustigmatophyte | *Nannochloropsis oculata* | SAG | 38.85 |
| green algae | *Ankistrodesmus falcatus* | SAG | 202-2 |
| green algae | *Botryococcus braunii* | SAG | 30.81 |
| green algae | *Botryococcus braunii* | CCAP | 807/1 |
| green algae | *Botryococcus braunii* | CCAP | 807/2 |
| green algae | *Botryococcus braunii* | NIES | 2199 |
| green algae | *Botryococcus braunii* | Eawag isolate | PT1 |
| green algae | *Botryococcus braunii* | N/A | Showa (US) |
| green algae | *Chlamydomonas reinhardtii* | CC | 1690 |
| green algae | *Chlamydomonas reinhardtii* | CC | 3060 |
| green algae | *Chlorella vulgaris* | SAG | 211-11b |
| green algae | *Cosmarium botrytis* | SAG | 136.8 |
| green algae | *Desmodesmus abundans* | Eawag isolate | GM26 |
| green algae | *Desmodesmus armatus* | Eawag isolate | L t0 |
| green algae | *Kirchneriella subcapitata* | SAG | 12.81 |
| green algae | *Lagerheimia hindakii* | SAG | 11.92 |
| green algae | *Lagerheimia subsalsa* | Eawag isolate | GM37 |
| green algae | *Medakamo hakoo* | NIES | 4000 |
| green algae | *Messastrum gracile* | Eawag isolate | GM45 |
| green algae | *Micractinium pusillum* | CCAP | 231/1 |
| green algae | *Oocystella heteromucosa* | Eawag isolate | GM35 |
| green algae | *Oocystis solitaria* | SAG | 83.8 |
| green algae | *Oocystis* sp. | Eawag isolate | L t0 |
| green algae | *Pandorina unicocca* | Eawag isolate | GM17 |
| green algae | *Pediastrum boryanum* | SAG | 87.81 |
| green algae | *Pediastrum boryanum* | Eawag isolate | GM20 |
| green algae | *Pediastrum duplex* | SAG | 261-2 |
| green algae | *Pediastrum duplex* | Eawag isolate | GM22 |
| green algae | *Scenedesmus acuminatus* | SAG | 38.81 |
| green algae | *Scenedesmus armatus* | Eawag isolate | GM28 |
| green algae | *Scenedesmus ellipticus* | Eawag isolate | GM25 |

To help overcome these knowledge gaps, we conducted a systematic screening of 49 algae strains, comprising 40 species, from taxonomically diverse groups of phytoplankton in order to broaden our understanding of high pH/alkalinity tolerance in algae. This bioprospecting effort focuses specifically on freshwater microalgae because their tolerance to high pH and alkalinity is largely unknown at present. The two main goals of this effort are (1) to identify novel algae strains capable of growing in high pH and high alkalinity conditions which could be used in commercial farming, and (2) to test whether a set of reportedly harmful algae species can also grow in high pH and high alkalinity conditions. We expected only a fraction of the strains tested to be productive in these extreme conditions but also expected to discover additional alkaline-tolerant strains which could be of value for industrial algae cultivation for direct air capture of carbon dioxide into value-added bioproducts.

## Materials and Methods

### Algae cultures

The Aquatic Ecology Department of Eawag maintains a large collection of microalgal strains representing a broad array of taxa from unique environmental origins. This collection presented a unique opportunity to bioprospect for high pH/alkalinity tolerance across much of the freshwater phytoplankton tree of life. The cultures selected for this experiment range from recent isolates from Swiss lakes to many standard “type strains” obtained from culture collections. This allows us to test not only lab-adapted model strains but also those more recently isolated from natural settings. With the exception of a few brackish strains (e.g., *Nannochloropsis oculata*), the microalgae used are assumed to be adapted primarily to freshwater (i.e., this study does not include strictly marine strains). The strains used in the experiment are listed in Table 1. Before the experiment, cultures were maintained at 20 °C in COMBO medium with double the concentration of all nutrients described in the original medium recipe^26^; this medium is hereafter referred to as 2X COMBO. This common medium was used for all strains, as it is designed to allow growth of many distinct algal taxa (e.g., diatoms, green algae, cyanobacteria); in terms of major nutrients, 2X COMBO contains 2000 µM N, 100 µM P, and 200 µM Si (see Kilham et al.^26^ for full recipe).

### Experimental design

The primary factor that was experimentally manipulated in this experiment (apart from algal strain identity) was the medium type. The study is organized into three phases, with each phase exposing algae to a growth medium with a higher level of pH/alkalinity (as well as salinity) than the previous one. The rationale for this phased approach was that stepwise acclimation may reveal tolerances that would not be observed by directly subjecting algae to more extreme conditions. In each phase, the three different media types tested were: (1) 2X COMBO, which served as a positive control; (2) 2X COMBO with added NaCl at increasing levels from phase to phase, which served to distinguish salinity tolerance from pH/alkalinity tolerance; and (3) 2X COMBO with added NaHCO_3_ and/or Na_2_CO_3_ to identify high pH/alkalinity tolerance. The salinity treatments match the mass concentration of salinity contributed by the added carbonate salts; this results in a 44% higher molar concentration of NaCl versus carbonate salts in each phase (e.g., 2.1 g/L of NaCl equals 35.9 mM, while 2.1 g/L of NaHCO3 equals 25 mM). Table 2 describes the media treatment levels used for each of the three phases. In Phase 1, only NaHCO_3_ was used to add alkalinity (and moderately increase the initial pH to near 8.5); in Phases 2 and 3 an equimolar mixture of NaHCO_3_ and Na_2_CO_3_ was used as this creates a buffered system with an initial pH near 10. The total alkalinity increase due to these sources is described as: *carbonate alkalinity = [HCO_3_ ^-^] + 2×[CO_3_ ^2-^]*.

**Table 2.** Experimental treatments used in each phase.

| Phase | Control treatment | NaCl treatment | High pH & alkalinity treatment |
| --- | --- | --- | --- |
| **Phase 1:**<br>**acclimation** | 2X<br>COMBO (pH 7.5) | 2.1 g/L NaCl<br>(pH 7.5) | 2.1 g/L NaHCO<sub>3</sub><br>(pH 8.49; 25 mM carbonate alkalinity) |
| **Phase 2:**<br>**high alkalinity**<br>**screening** | 2X<br>COMBO<br>(pH 7.5) | 4.75 g/L NaCl<br>(pH 7.5) | 2.1 g/L NaHCO<sub>3</sub> and 2.65 g/L Na<sub>2</sub>CO<sub>3</sub><br>(pH 10.09; 75 mM carbonate alkalinity) |
| **Phase 3:**<br>**extreme alkalinity**<br>**screening** | 2X<br>COMBO<br>(pH 7.5) | 9.5 g/L NaCl<br>(pH 7.5) | 4.20 g/L NaHCO<sub>3</sub> and 5.30 g/L Na<sub>2</sub>CO<sub>3</sub><br>(pH 10.02; 150 mM carbonate alkalinity) |

With the exception of the above media manipulations, each phase used identical methods for cultivation of microalgae. Specifically, 900 µL of medium per well was dispensed into sterile 48-well plates and 100 µL algae culture was added to yield a total volume of 1 mL per well, with 4 replicates per treatment. The phased acclimation approach was carried out as follows: for Phase 1, the 100 µL algal inoculum came from 50 mL flask pre-cultures; for Phase 2, the 100 µL algae came directly from corresponding media treatment wells at the end of Phase 1; similarly, the 100 µL algae to start Phase 3 came directly from corresponding Phase 2 wells. This was done to ensure that the algae populations acclimated to a given phase would be directly tested for their tolerance to the subsequent and progressively harsher environment. It also mimics, in a microcosm environment, the act of transferring algae cultures via a 10% v/v dilution, as might be done when scaling up commercial cultivation from smaller to larger raceway volumes, and it maintains a constant volume proportion of new medium across treatments. A potential limitation of this sequential transfer approach is that while the wells were always inoculated with the same volume of algae culture (100 µL; 10% by volume), they were not equalized in terms of initial algal biomass. Therefore, the reader should be aware that differences in initial densities across both strains and media treatments could influence the resulting growth parameters. The duration of experimental phases were 7 days (Phase 1); 15 days (Phase 2); and 10 days (Phase 3). The duration of Phase 1 was chosen to provide a week-long acclimation period, while the durations of Phase 2 and 3 were chosen to allow all cultures to reach carrying capacity in a given treatment. In all phases, algae were grown in a Memmert ICP 700 incubator set to 24 °C with cool white fluorescent lights set to 14h:10h light:dark cycle and 98.2 ± 14.3 SD µmol photons m^-2^ s^-1^. All plates were covered with Breathe Easy^®^ membranes (Diversified Biotech) to allow for gas exchange and minimize evaporation. Growth was tracked using *in vivo* fluorescence (recorded in terms of relative fluorescence units, RFU) as a proxy for total algal biomass. Chlorophyll-a fluorescence was measured for all experimental units with excitation/emission at 460/685 nm, using a Biotek® Cytation 5 plate reader. Additionally, phycocyanin fluorescence was used as a proxy to track growth of cyanobacteria with excitation/emission wavelengths of 586/647 nm. Fluorescence was chosen over optical density as a biomass proxy because it is more specific to photosynthetically active algal biomass (i.e., it reduces the effect of changes in non-photosynthetic bacterial abundance, cellular debris, or mineral precipitates); however, the reader should note that, as with any biomass proxy, fluorescence should not be interpreted as a perfect substitute for direct biomass measurements (e.g., ash free dry weight). Measuring ash free dry weight becomes logistically prohibitive for high-throughput screening studies such as this.

### Statistical analysis

The two main metrics used to assess treatment effects in this study are (1) the maximum fluorescence attained over the course of an experimental phase, which serves as a proxy for maximum algal biomass and (2) the growth rate over time. Each of these values were calculated for each experimental unit (n = 4 replicates for each combination of strain/treatment/phase). It should again be noted that max. fluorescence can be affected by both algal biomass and per-cell chlorophyll, thus representing a relative (and not absolute) proxy of peak photosynthetically active biomass, and that initial densities in each phase may affect both the growth rate and maximum biomass. Treatment effects on max. fluorescence are shown in figures in terms of the mean % change effect caused by a treatment relative to the average max. fluorescence of the control, as well as the 95% confidence intervals of the effect. All analyses were performed in R version 4.4.3^27^. All growth curves used to calculate summary growth data are provided in the Supporting Information (Fig. S1, S2, S3).

We assessed several complementary methods for estimating growth rates to ensure that the maximum specific growth rates obtained were realistic and not biased by a particular model fit. Specifically, we tested exponential, logistic, and Gompertz model fits using v0.8.4 of the R package ‘growthrates’^28^, a suite of models in v1.2 of the R package ‘growthTools’^29^ in which the best of five models applied is selected by AICc scores, as well as the “manual approach” of calculating growth using the formula *µmax =* [*ln(fluorescence day 4/fluorescence day 0)]/(4 days)*; this period was chosen as the first four days best captured exponential growth when considering all strains and environments. The different methods yielded similar median growth rate estimates (Fig. S4), and growth rates tended to be highly correlated (Pearson’s *r* up to 0.96), with the exception of the logistic and Gompertz models, which often overestimated growth, resulting in many outliers with unrealistic growth rate estimates (Fig S4, S5), making these models overall inappropriate. Overall, we find that the choice of growth rate estimation method among the three remaining suitable methods does not substantially affect inferences drawn from our data. In the main text, we show growth rate estimates obtained using the ‘growthTools’ package, as this method allows for incorporation of lag phases (observed in ca. 8% of cases), which yields a slightly more accurate estimate of exponential growth after a lag compared to the other options, and has the highest mean model R^2^ value (Fig. S6).

As initial densities varied substantially in the acclimation period (Phase 1), we do not assess growth rates during this time, as these density dependent effects likely influenced growth rates. However, initial densities for Phases 2 and 3 are meant to represent approximately 10% of carrying capacity for each strain × environment combination (due to 10% v/v transfer of stationary phase cultures), which makes growth rate comparisons across strains and treatments more appropriate. Effects of initial densities are explored further in Fig. S7; in summary, we observe no overall significant effect of variation in initial density on growth rates in Phase 2, although the effects of initial densities differ among the media types (i.e., increasing initial density had no effect for the pH & alkalinity treatments but a slightly negative effect for controls).

Welch’s ANOVAs were used to test for significant differences among the three treatments for each strain in each phase, as these allow for heterogeneity of variances, which was observed throughout the data. Games Howell post hoc tests (analogous to Tukey tests) were performed to test for differences between means as they also allow for heterogeneous variances and control for type 1 error. We tested the ability of salinity responses to predict alkalinity responses using a combination of linear models and Spearman correlation tests.

## Results

### Phase 1: acclimation to moderate alkalinity conditions

In the first experimental phase, the addition of alkalinity as NaHCO_3_ had a generally positive effect on algal biomass (Fig. 1). Twenty-one of the 49 strains (43%) reached a significantly higher maximum fluorescence value than their respective controls (see Table S1 for Games Howell test statistics); moreover, 19 of these strains had a 25% or greater increase in biomass relative to controls and 4 strains had a biomass increase of over 100% with NaHCO_3_ addition. Of the remaining strains, 15/49 (30.6%) showed no significant effects of NaHCO_3_ addition, while 13/49 (26.5%) had significantly reduced biomass in terms of *in vivo* fluorescence under the moderate increase in pH/alkalinity (25 mM carbonate alkalinity).

**Fig. 1.**
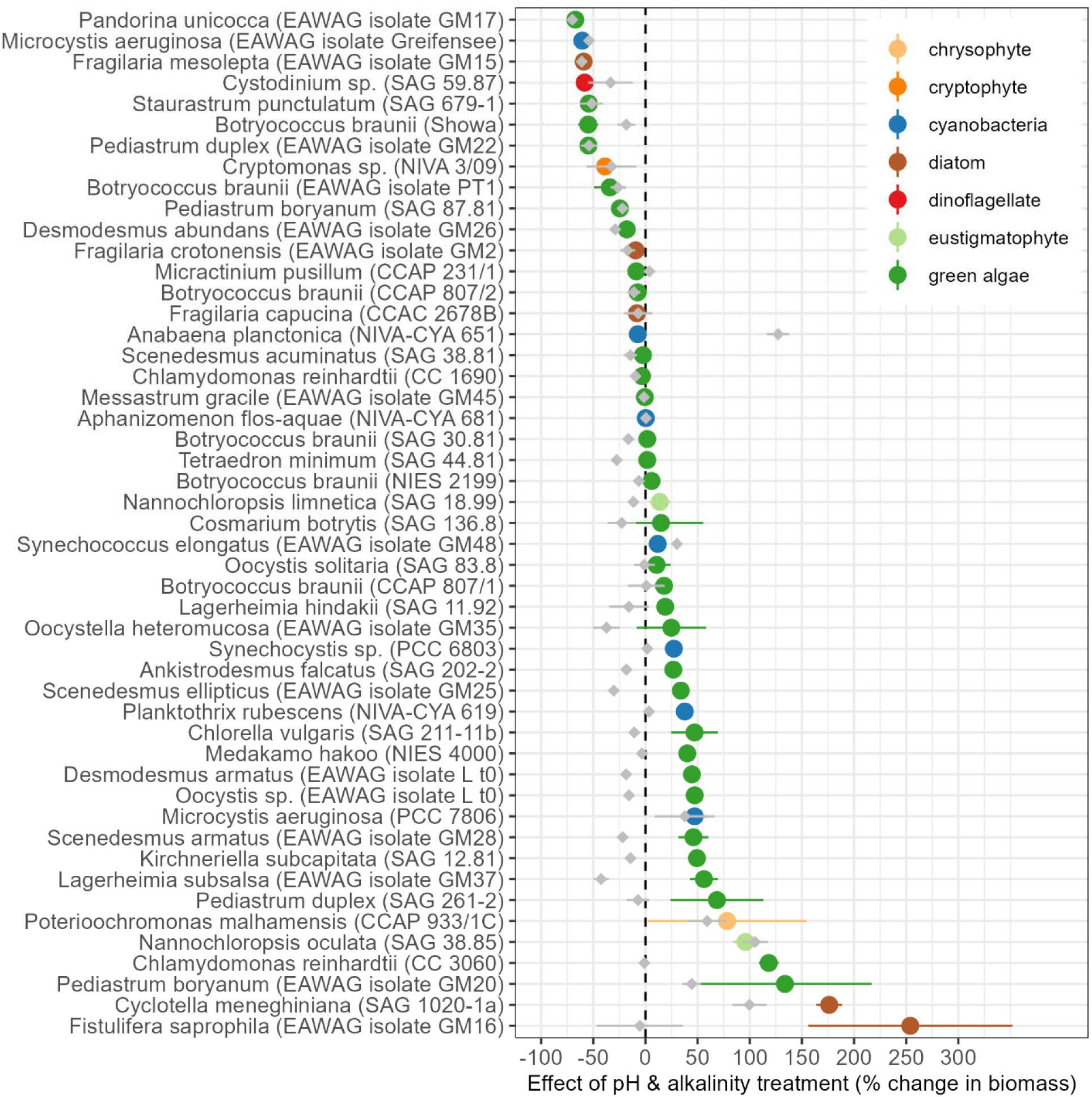
Effects of the pH and alkalinity treatment on algal biomass during Phase 1 (i.e., during the 7-day acclimation phase to a mild increase in initial pH and alkalinity: pH 8.5 and 25 mM of alkalinity added as carbonates). Colored circles and corresponding lines represent the mean and 95% confidence interval of the pH/alkalinity treatment effect in the form of maximum biomass production estimated as *Effect size = (treatment max. fluorescence - mean control max. fluorescence)/(mean control max. fluorescence)*. Gray diamonds and lines show the mean and 95% CI effect of salinity (2.1 g/L NaCl during Phase 1) for each strain. Values above zero indicate a positive effect of the treatment relative to control, zero indicates no effect, and negative values indicate a detrimental effect of the pH/alkalinity (or salinity) treatment.

Compared to effects of pH/alkalinity, the magnitude of salinity treatment (2.1 g/L NaCl) effects on algal growth was relatively low (i.e., gray diamonds in Fig. 1), especially when compared to algae strains with highly positive responses to pH/alkalinity. Although the effects of NaCl addition at 2.1 g/L were weaker overall than the effects of 2.1 g/L NaHCO_3_ addition, there was a tendency towards negative effects, with 23/49 strains significantly inhibited in growth and only 6/49 showing higher growth with NaCl versus the control (Table S1). While several strains had positive responses to both NaHCO_3_ and NaCl of roughly equal magnitude (e.g., *Nannochloropsis oculata* and *Cyclotella meneghiniana*), many others exhibited drastically different responses to the two treatments, and in the case of *Anabaena planctonica* a strong positive response to NaCl but no effect of NaHCO_3_. In total, 25/49 strains had significantly higher max. fluorescence with alkalinity than salinity, 19/49 had equivalent effects of each, and only 5 had higher max. fluorescence with added salinity than added alkalinity. A linear regression indeed shows that while NaCl responses are significantly related to NaHCO_3_ responses, they only explain 24% of the variance in NaHCO_3_ response effects (*F*_1,47_ = 16.4, *p* = 0.0002). In other words, data from this portion of the experiment show that algal responses to moderately alkaline conditions tend to be decoupled in magnitude from their responses to moderately saline conditions. These results also show that moderate bicarbonate addition often (but not always) has a significant growth stimulating effect which can be observed across a suite of taxonomically distinct algae such as chrysophytes, cyanobacteria, eustigmatophytes, green algae, and diatoms.

While there is a clear growth enhancement of several industrially relevant production strains (e.g., *Nannochloropsis oculata*, *Chlorella vulgaris,* and *Chlamydomonas reinhardtii*) in Phase 1, there is also a growth enhancement of certain potentially toxic cyanobacteria (e.g., *Planktothrix rubescens* and *Microcystis aeruginosa*) as well as the mixotrophic grazer *Poterioochromonas malhamensis*, which is a widespread and destructive pest in algae farms^25^. There is also evidence for within-species variation in tolerance to high pH/alkalinity; the 6 unique strains of the hydrocarbon-rich alga *Botryococcus braunii* which we tested in this study have distinct responses to increased pH/alkalinity, ranging from minor increases (e.g., strain CCAP 807/1) to drastic decreases (e.g., strain Showa) in growth (Fig 1, Fig. S1). Additional evidence exists for within-species differences in responses, e.g., the two unique strains of *Pediastrum duplex* have diverging responses to the alkalinity treatment in Phase 1, as do the two strains of *Pediastrum boryanum*. Within-genus differences are also noteworthy in, e.g., *Desmodesmus*, *Scenedesmus*, and *Nannochloropsis*.

### Phase 2: screening for tolerance to high pH and alkalinity levels

In contrast to the observed results for Phase 1, nearly all strains (47/49) had reduced maximum biomass compared to the controls when subjected to high pH and alkalinity in Phase 2 (15 days of growth at pH 10 and 75 mM of carbonate alkalinity; inoculated with a 10% v/v transfer from Phase 1 cultures, Fig. 2). Only *Synechocystis* PCC 6803 had greater biomass in the high pH/alkalinity treatment than in the control, while all but one strain (with no significant effect) had significant decreases (Table S2). Analysis of growth rates shows that approximately 16% of the strains (8/49) had positive average growth rates under these conditions (Fig. 3). The consistent differences between the salinity and the pH/alkalinity treatments on both biomass and on growth rates again indicate that, in most cases, the impact on growth is likely due to pH/alkalinity *per se* rather than due to the increase in salinity that is concomitant with increased alkalinity. For a relatively small subset of strains, however, we do observe very similar responses to salinity and pH/alkalinity (see, e.g., *Pandorina unicocca*, *Fragilaria capucina*, and *Staurastrum punctulatum*). In contrast to Phase 1, all potentially toxic or harmful “pest” species were severely inhibited and were incapable of growth in the high pH/alkalinity conditions of Phase 2. Instead, species with positive growth rates included industrially important strains of green algae from the family Scenedesmaceae (*Desmodesmus* and *Scenedesmus*) and the cyanobacteria (*Synechocystis* and *Synechococcus*).

**Fig. 2.**
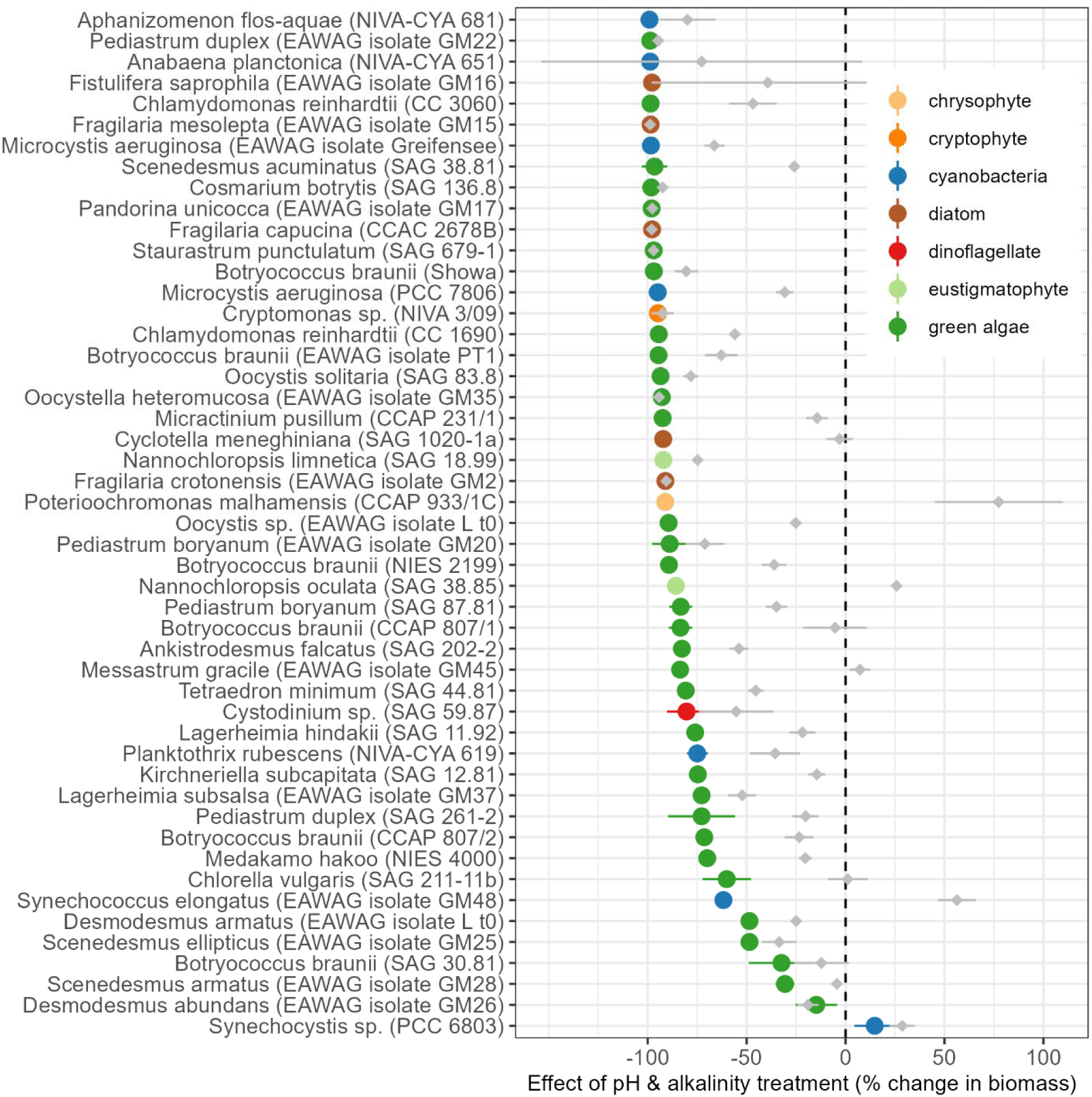
Effects of the pH and alkalinity treatment on algal biomass during Phase 2 (i.e., the 15-day screening in high pH and alkalinity – pH 10 and 75 mM of alkalinity added as carbonates). Colored circles and corresponding lines represent the mean and 95% confidence interval of the pH/alkalinity treatment effect in the form of maximum biomass production estimated as *Effect size = (treatment max. fluorescence - mean control max. fluorescence)/(mean control max. fluorescence)*. Gray diamonds and lines show the mean and 95% CI effect of salinity (4.75 g/L NaCl during Phase 2) for each strain. Values above zero indicate a positive effect of the treatment relative to control, zero indicates no effect, and negative values indicate a detrimental effect of the pH/alkalinity (or salinity) treatment.

**Fig. 3.**
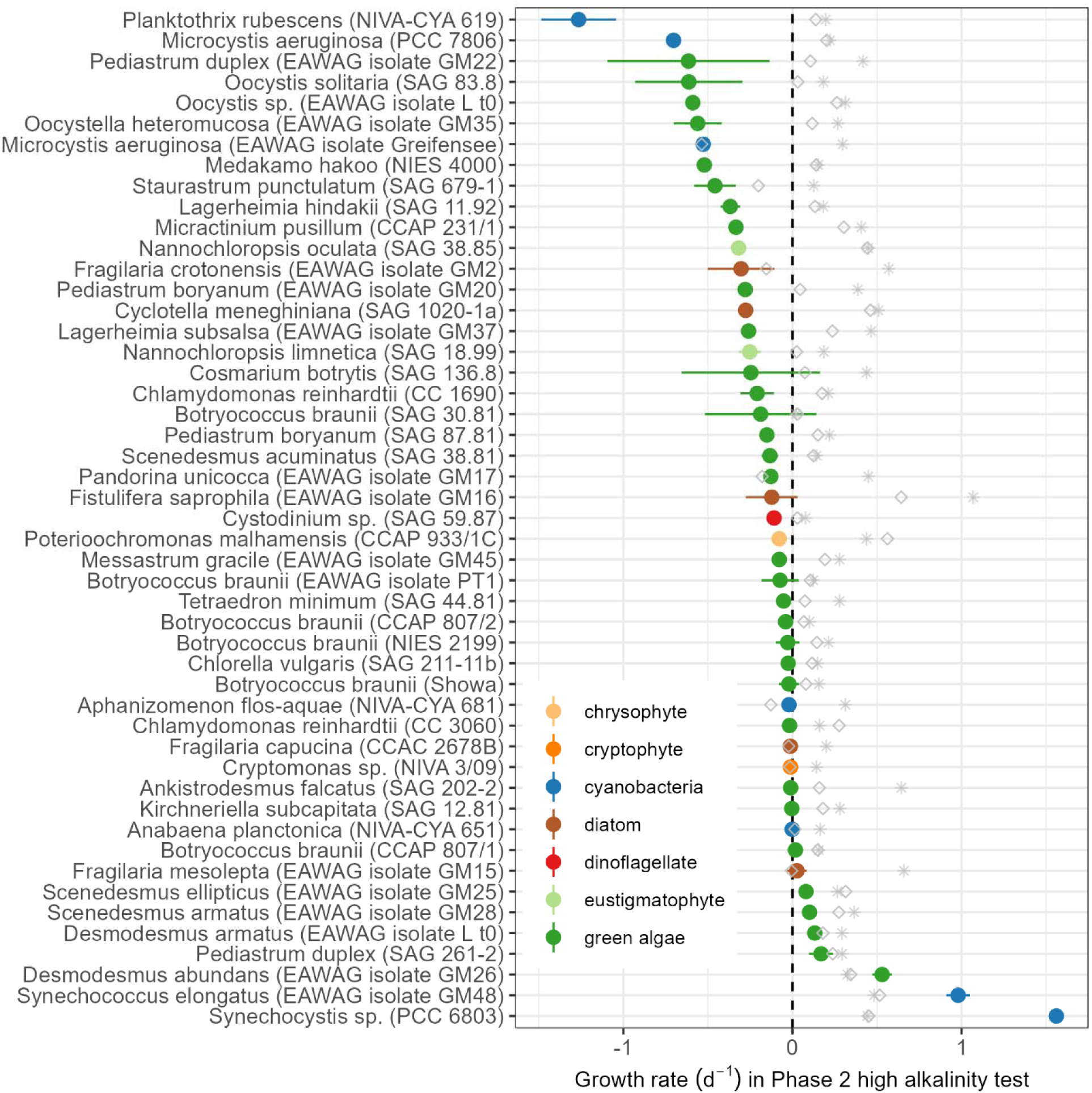
Growth rates of each algal strain in the high pH and alkalinity screening (Phase 2, 15-day screening at pH 10 and 75 mM of alkalinity added as carbonates) estimated using the R package ‘growthTools’. Colored circles and corresponding lines represent the mean and 95% confidence interval of growth rate in high pH and alkalinity; gray diamonds show the average growth rates of the salinity treatments (4.75 g/L NaCl), and asterisks show the average growth rates in the control treatment.

### Phase 3: screening for tolerance to extreme pH and alkalinity levels

Only the 10 best performing strains in Phase 2 were transferred to the extreme conditions of Phase 3. The extreme pH and alkalinity levels in Phase 3 reduced biomass by over 80% for all strains (Fig. 4), and only one strain (*Desmodesmus abundans*) was able to maintain a positive growth rate (Fig. 5). Again, the effects of pH/alkalinity and salinity were decoupled; although salinity generally had a negative effect compared to the control, it was much weaker than the pH/alkalinity effect. In the case of *Synechocystis* PCC 6803, salinity appeared to induce a mild increase in maximum biomass.

**Fig. 4.**
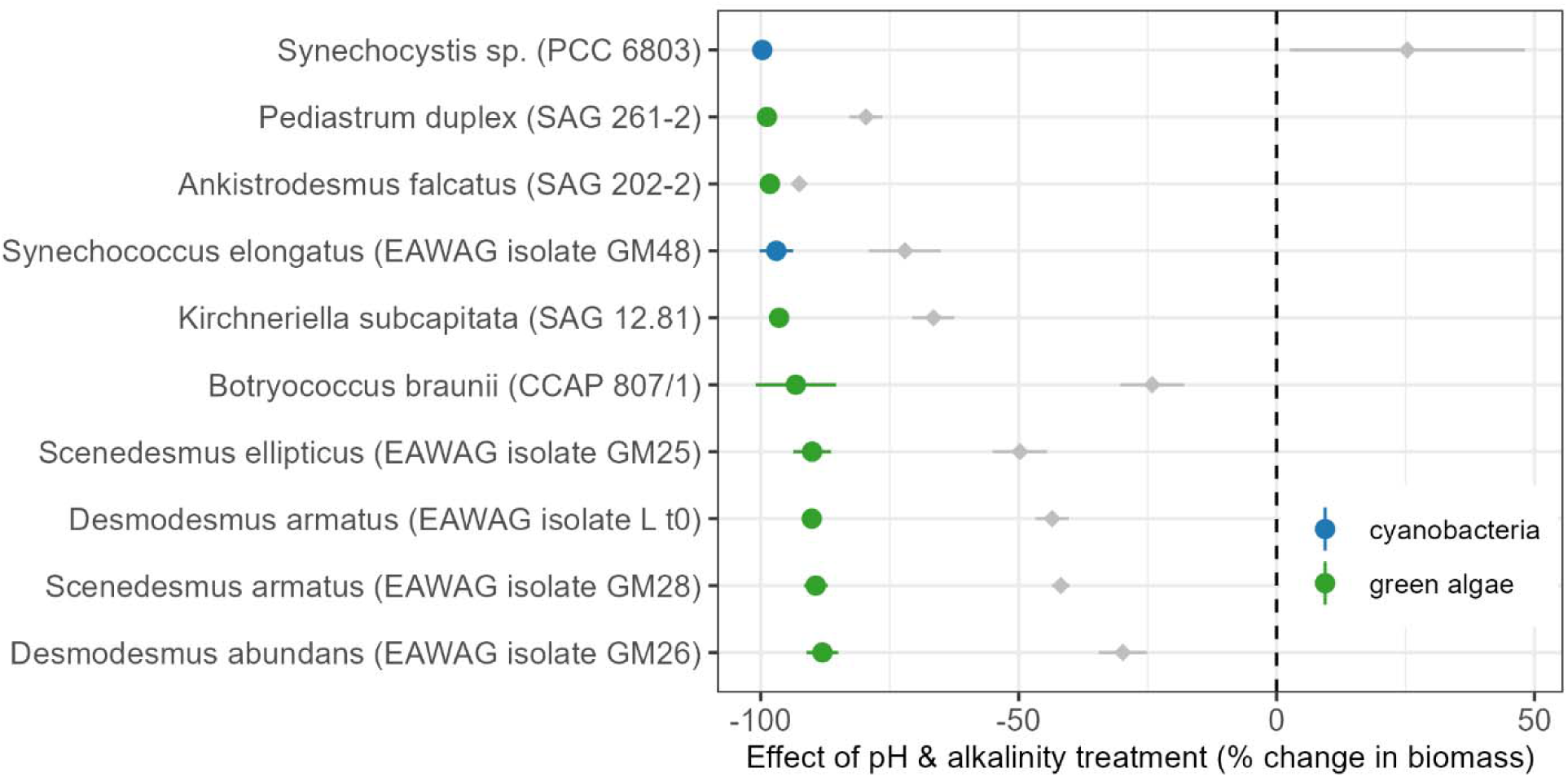
Effects of the pH and alkalinity treatment on algal biomass during Phase 3 (i.e., the 10-day screening in extreme pH and alkalinity – pH 10 and 150 mM of alkalinity added as carbonates). Colored circles and corresponding lines represent the mean and 95% confidence interval of the pH/alkalinity treatment effect in the form of maximum biomass production estimated as *Effect size = (treatment max. fluorescence - mean control max. fluorescence)/(mean control max. fluorescence)*. Gray diamonds and lines show the mean and 95% CI effect of salinity (9.5 g/L NaCl during Phase 3) for each strain. Values above zero indicate a positive effect of the treatment relative to control, zero indicates no effect, and negative values indicate a detrimental effect of the pH/alkalinity (or salinity) treatment.

**Fig. 5.**
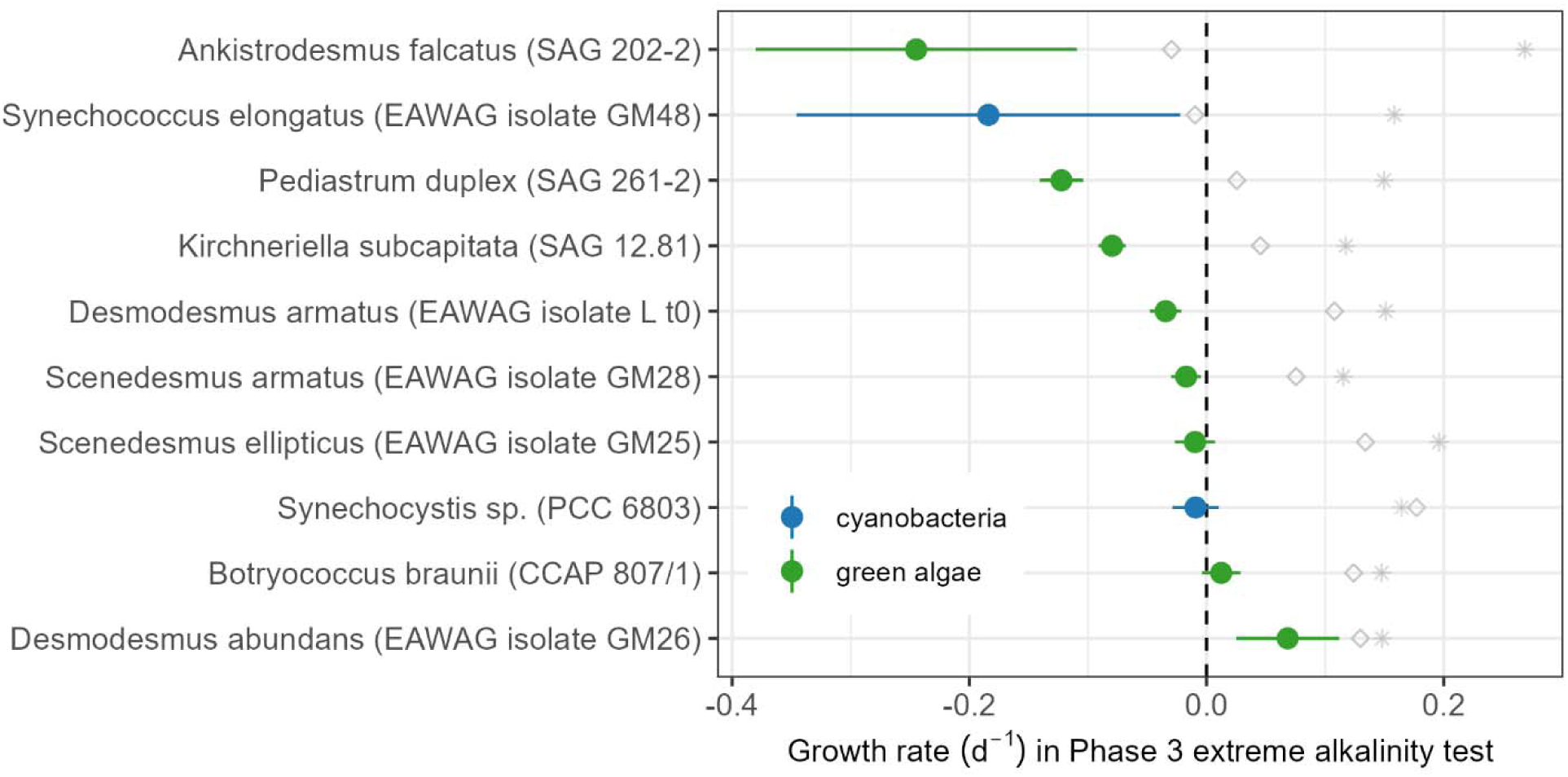
Growth rates of each algae strain in the high pH and alkalinity screening (Phase 3), estimated using the R package ‘growthTools’. Filled circles and lines represent the mean and 95% confidence interval of growth rate in high pH and alkalinity; gray diamonds show the average growth rates the salinity treatment (9.5 g/L NaCl), and asterisks show the average growth rates in the control treatment.

### Alkalinity effects are not driven by salinity stress

Increased alkalinity inherently increases salinity in aquatic systems by raising the concentration of dissolved ionic solutes, creating a link between the two factors that often makes causal and mechanistic inference difficult. We do observe an overall significant but weak correlation between salinity and alkalinity effects (Pearson’s *r* = 0.50, p < 0.01). However, our results across all three phases also show that salinity effects can differ substantially from alkalinity effects; Fig. 6A shows the quantitative relationship between the pH/alkalinity and salinity treatment effects on biomass production in all three phases, and that in all phases the alkalinity-salinity relationship is never one-to-one, as would be expected if salinity stress was driving responses to high alkalinity treatments. Similarly, the relationship between high salinity growth rates and high pH/alkalinity growth rates nearly always deviates from the 1:1 line, with most points showing that growth is positive in high salinity but negative for the same strain when subjected to alkalinity at the same concentration (Fig. 6B).

**Fig. 6.**
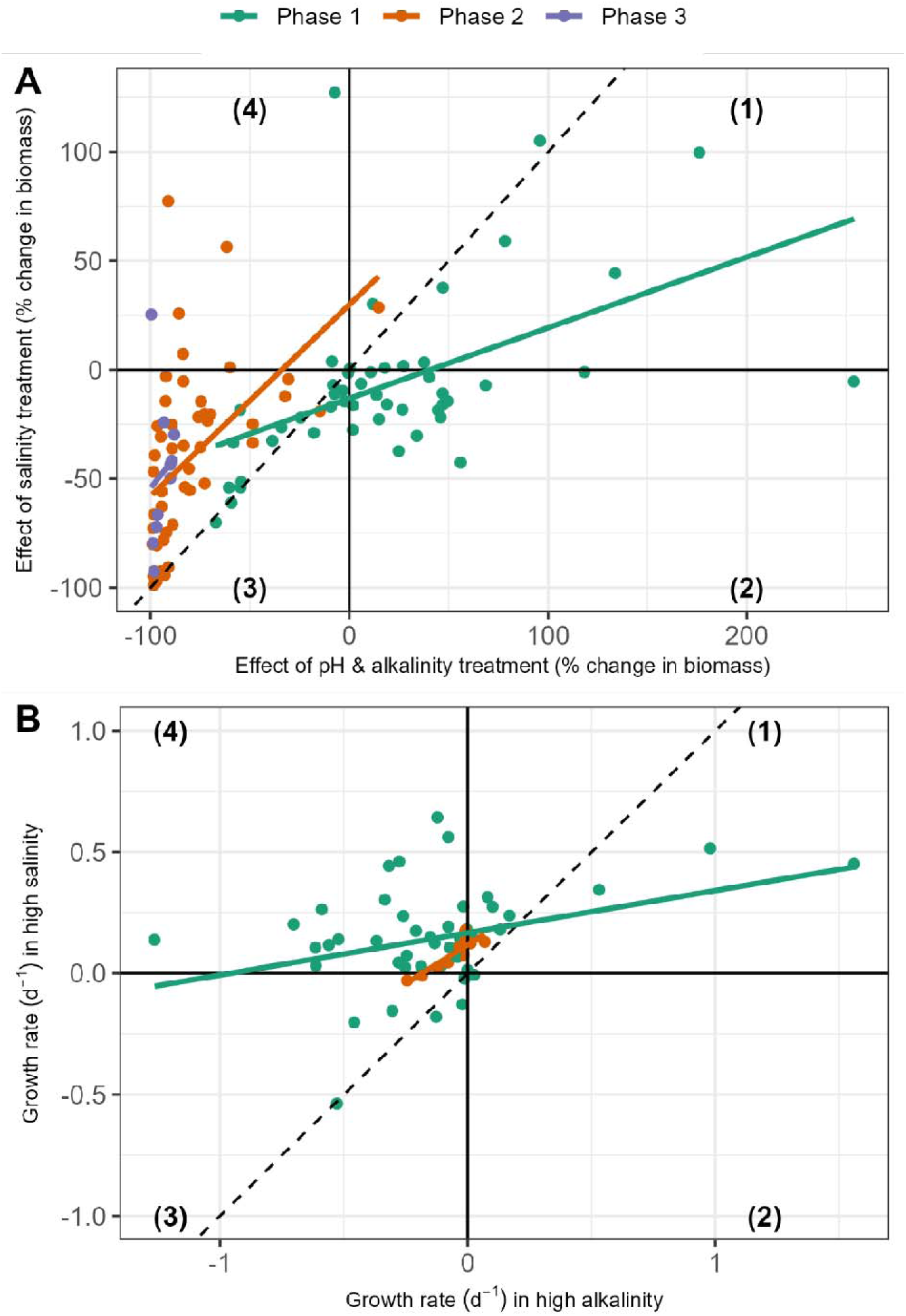
Relationship between pH/alkalinity effect size and salinity effect size on max. fluorescence (A), and relationship between growth rate in pH/alkalinity versus salinity treatments (B), grouped by each experimental phase. As in previous figures, effects on biomass are calculated as *Effect size = (treatment max. fluorescence - mean control max. fluorescence)/(mean control max. fluorescence)*. Effect sizes (A) and growth rates (B) can each be split into four quadrants: (1) positive for both treatments; (2) positive for alkalinity but negative for salinity; (3) negative for both; and (4) negative for alkalinity but positive for salinity. In Phase 1, the decoupling of alkalinity and salinity results in more positive effects of alkalinity, while the inverse is true for the more extreme conditions in Phases 2 and 3. Points show means for each algal strain and phase, solid lines show linear model fits by phase, and the dashed lines show a 1- to-1 relationship which would indicate equal effects of salinity and alkalinity.

## Discussion

This study represents the largest and taxonomically broadest screening of algal tolerance to high pH and alkalinity to date to our knowledge and provides much-needed information on the innate ability of freshwater algae to tolerate conditions that are generally considered suitable only for the growth of extremophiles. Our study shows that the magnitude of pH and alkalinity stress significantly alters the suite of species capable of productive growth. The relatively low pH/alkalinity in Phase 1 (∼pH 8.5 and 25 mM carbonate alkalinity) yielded growth increases for many algae (including both desired and undesired strains), while increasing the pH to 10 and further increasing the alkalinity in Phase 2 and 3 inhibited the majority of strains, again reinforcing the strong selective force of such extreme pH and alkalinity conditions. Additionally, our study begins to disentangle the effects of alkalinity versus salinity, which are generally co-varying and, thus, sometimes confounding factors in similar studies. Here we clearly show that salinity and alkalinity effects are often decoupled across algal groups, as only a small subset of strains appeared to have similar responses to both high salinity and high alkalinity growth conditions.

Our study is distinct from previous ones because our high-throughput screening provides relatively broad taxonomic information about tolerance to alkalinity and salinity; however, this broad view prevents an in-depth analysis of each individual strain, as has been the focus of previous studies. Specifically, earlier studies showed how lipid content and other important biochemical characteristics were affected by increased alkalinity, usually with significant increases in lipid accumulation^30–32^. Our screening does not assess biomass quality. However, our study does clearly show that the beneficial impacts of moderate added alkalinity (25 mM) on industrial strains could also stimulate the growth of undesired or potentially harmful taxa. Specifically, we observed that the grazer *Poterioochromonas* and the toxic bloom-forming algae *Microcystis* and *Planktothrix* all had increased growth at 25 mM added alkalinity relative to the control (Fig. 1, Fig. S1). Therefore, while we know from previous work that moderate increases in alkalinity can serve to increase lipid accumulation, our study suggests that such levels may be insufficient to yield the co-benefit of acting as a crop protection measure. On the other hand, we observed inhibition of all potentially harmful species in our study with higher levels of pH (∼10) and alkalinity (75 mM). Another caveat of our results is that the growth rates we observed would likely differ with different growth conditions (e.g., light, temperature, nutrients) and that the different initial densities due to the serial transfer experimental design may have introduced some variability in the exact growth rate observed (Fig. S7). However, the main outcome of importance is whether a certain strain was able to achieve positive growth in a certain environment, and our results definitively show which strains are (or are not) capable of growth in each high pH/alkalinity environment.

While our study confirms that extremophiles are indeed rare, it does suggest that the potential of microalgae from relatively “normal” aquatic systems should not be ignored. Previous work has employed bioprospecting efforts and identified productive alkaliphilic algae from alkaline lakes such as Soap Lake in Washington^19,23^. We suggest that alkaline systems are a clear and logical venue for bioprospecting, and that this should continue; however, there may also be hidden potential in your local puddle, as common pond algae like *Scenedesmus* and its relatives, as well as common cyanobacteria, were the most successful in our high alkalinity screening. A key benefit of bioprospecting for locally sourced alkaliphilic strains is that they are likely to be better adapted to local climate conditions, whereas strains from a select few geographically/climatically distant alkaline lakes may not have traits conferring optimal adaptation to local conditions. Future work may also benefit from combining the high alkalinity approach with other strategies, e.g., using rationally designed polycultures of alkaline-tolerant strains and probiotic bacteria to stabilize outdoor production, or the use of adaptive laboratory evolution to push the limits of tolerance to high alkalinity and pH in commercial algae strains. Experimental gradient designs with more treatment levels (i.e., more than the three levels used here) will also further elucidate pH and alkalinity tolerance thresholds to optimize productivity in key algal species. A synergistic approach combining high pH and alkalinity cultivation with other advances in the ecology and engineering of algae farming will likely multiply the observed benefits of such strategies.

## Conclusions

Our study screened 49 algal strains (including green algae, cyanobacteria, diatoms, and others) for the ability to grow in three increasingly harsh levels of pH and alkalinity. We show that moderate increases in pH and alkalinity (initial pH of 8.5 and 25 mM added carbonate alkalinity) cause significant growth increases in many strains; however, these include both desired production strains as well as potentially harmful algae. High alkalinity levels (initial pH of 10 and 75 mM carbonate alkalinity) broadly inhibited growth of most species, with only 8/49 strains showing positive average growth rates, and only one species growing in the extreme condition (initial pH of 10 and 150 mM carbonate alkalinity). This expands our knowledge of the species pool that we may expect to grow in commercial algae farms using such cultivation strategies and identifies key algal taxa that generally have the ability to be highly productive in alkaline cultivation systems at various levels of pH and alkalinity.

## Supporting Information

Supplementary Tables (tables with test statistics to assess treatment effects); Supplementary Figures (growth curves for all experimental units; figures analyzing different growth rate estimation methods, analysis of initial density effects).

## Acknowledgements

We would like to thank Gabriella Mege for isolating many of the strains in the Eawag culture collection; Shigeru Okada, Lev Typsin, and Sammy Pontrelli for providing additional algal cultures; Al Parker for help and guidance with statistical analysis; and Marta Reyes, Raphaël Bossart, and Silvana Käser for additional help with isolation, maintenance, and characterization of algae strains. This research was funded by a seed grant from Eawag and through the United States National Science Foundation under grant #2125083 (URoL-MIM).

## Data availability

All data and code used will be publicly available on Zenodo upon acceptance.

## Author contributions

All authors contributed to the conception and design the study; PKT conducted the experiment; PKT performed statistical analyses, data visualizations, and wrote the draft manuscript; RG and AN reviewed and edited the draft manuscript; all authors approve its final version for submission.

## References

(1) Olabi, A. G.; Shehata, N.; Sayed, E. T.; Rodriguez, C.; Anyanwu, R. C.; Russell, C.; Abdelkareem, M. A. Role of Microalgae in Achieving Sustainable Development Goals and Circular Economy. Sci. Total Environ. 2023, 854 (July 2022), 158689. 10.1016/j.scitotenv.2022.158689.

(2) Lane, T. W. Barriers to Microalgal Mass Cultivation. Curr. Opin. Biotechnol. 2022, 73, 323–328. 10.1016/j.copbio.2021.09.013.

(3) Chisti, Y. Constraints to Commercialization of Algal Fuels. J. Biotechnol. 2013, 167 (3), 201–214. 10.1016/j.jbiotec.2013.07.020.

(4) Day, J. G.; Gong, Y.; Hu, Q. Microzooplanktonic Grazers – A Potentially Devastating Threat to the Commercial Success of Microalgal Mass Culture. Algal Res. 2017, 27 (April), 356–365. 10.1016/j.algal.2017.08.024.

(5) Carney, L. T.; Lane, T. W. Parasites in Algae Mass Culture. Front. Microbiol. 2014, 5, 278. 10.3389/fmicb.2014.00278.

(6) Shurin, J. B.; Abbott, R. L.; Deal, M. S.; Kwan, G. T.; Litchman, E.; McBride, R. C.; Mandal, S.; Smith, V. H. Industrial-Strength Ecology: Trade-Offs and Opportunities in Algal Biofuel Production. Ecol. Lett. 2013, 16 (11), 1393–1404. 10.1111/ele.12176.

(7) Abdelaziz, A. E. M.; Leite, G. B.; Hallenbeck, P. C. Addressing the Challenges for Sustainable Production of Algal Biofuels: II. Harvesting and Conversion to Biofuels. Environ. Technol. 2013, 34 (13–14), 1807–1836. 10.1080/09593330.2013.831487.

(8) Pate, R.; Klise, G.; Wu, B. Resource Demand Implications for US Algae Biofuels Production Scale-Up. Appl. Energy 2011, 88 (10), 3377–3388. 10.1016/j.apenergy.2011.04.023.

(9) Singh, U.; Banerjee, S.; Hawkins, T. R. Implications of CO2 Sourcing on the Life-Cycle Greenhouse Gas Emissions and Costs of Algae Biofuels. ACS Sustain. Chem. Eng. 2023, 11 (39), 14435–14444. 10.1021/acssuschemeng.3c02082.

(10) Malavasi, V.; Soru, S.; Cao, G. Extremophile Microalgae: The Potential for Biotechnological Application. J. Phycol. 2020, 56 (3), 559–573. 10.1111/jpy.12965.

(11) Van Ginkel, S. W.; El-Sayed, W. M. M.; Johnston, R.; Narode, A.; Lee, H. J.; Bhargava, A.; Snell, T.; Chen, Y. Prevention of Algaculture Contamination Using Pesticides for Biofuel Production. Algal Res. 2020, 50 (December 2019), 101975. 10.1016/j.algal.2020.101975.

(12) Miller, I. R.; Bui, H.; Wood, J. B.; Fields, M. W.; Gerlach, R. Understanding Phycosomal Dynamics to Improve Industrial Microalgae Cultivation. Trends Biotechnol. 2024, 42 (6), 680–698. 10.1016/j.tibtech.2023.12.003.

(13) Fisher, C. L.; Lane, P. D.; Sale, K.; Lane, T. W. Persistence of Bacterial-Mediated Anti-Rotifer Protection in Preliminary Outdoor Cultivation Trial for Microchloropsis Salina. Algal Res. 2025, 86 (November 2024), 103942. 10.1016/j.algal.2025.103942.

(14) Newby, D. T.; Mathews, T. J.; Pate, R. C.; Huesemann, M. H.; Lane, T. W.; Wahlen, B. D.; Mandal, S.; Engler, R. K.; Feris, K. P.; Shurin, J. B. Assessing the Potential of Polyculture to Accelerate Algal Biofuel Production. Algal Res. 2016, 19, 264–277. 10.1016/j.algal.2016.09.004.

(15) Smith, V. H.; McBride, R. C.; Shurin, J. B.; Bever, J. D.; Crews, T. E.; Tilman, G. D. Crop Diversification Can Contribute to Disease Risk Control in Sustainable Biofuels Production. Front. Ecol. Environ. 2015, 13 (10), 561–567. 10.1890/150094.

(16) Yang, Y.; Tang, S.; Chen, J. P. Carbon Capture and Utilization by Algae with High Concentration CO2 or Bicarbonate as Carbon Source. Sci. Total Environ. 2024, 918 (January), 170325. 10.1016/j.scitotenv.2024.170325.

(17) Arora, N.; Tripathi, S.; Philippidis, G. P.; Kumar, S. Thriving in Extremes: Harnessing the Potential of PH-Resilient Algal Strains for Enhanced Productivity and Stability. *Environ*. Sci. Adv. 2025. 10.1039/D4VA00247D.

(18) Vadlamani, A.; Pendyala, B.; Viamajala, S.; Varanasi, S. High Productivity Cultivation of Microalgae without Concentrated CO2 Input. ACS Sustain. Chem. Eng. 2019, 7 (2), 1933–1943. 10.1021/acssuschemeng.8b04094.

(19) Vadlamani, A.; Viamajala, S.; Pendyala, B.; Varanasi, S. Cultivation of Microalgae at Extreme Alkaline PH Conditions: A Novel Approach for Biofuel Production. ACS Sustain. Chem. Eng. 2017, 5 (8), 7284–7294. 10.1021/acssuschemeng.7b01534.

(20) Chi, Z.; Elloy, F.; Xie, Y.; Hu, Y.; Chen, S. Selection of Microalgae and Cyanobacteria Strains for Bicarbonate-Based Integrated Carbon Capture and Algae Production System. Appl. Biochem. Biotechnol. 2014, 172 (1), 447–457. 10.1007/s12010-013-0515-5.

(21) Burch, T. A.; Hill, E. A.; Tamburro, J. M.; Bohutskyi, P.; Dennis, G.; Ashford, A.; LaPanse, A. J.; Beliaev, A.; Traller, J. C.; Pinowska, A.; Posewitz, M. C. A Selection Approach Using High oxygen and Pond-Mimicking Culture Conditions Increases Biomass Productivity in the Industrially Relevant Diatom Nitzschia Inconspicua Str. Hildebrandi. Algal Res. 2024, 77 (November 2023), 103333. 10.1016/j.algal.2023.103333.

(22) Ciferri, O. Spirulina, the Edible Microorganism. Microbiol. Rev. 1983, 47 (4), 551–578. 10.1128/mmbr.47.4.551-578.1983.

(23) Gao, S.; Pittman, K.; Edmundson, S.; Huesemann, M.; Greer, M.; Louie, W.; Chen, P.; Nobles, D.; Benemann, J.; Crowe, B. A Newly Isolated Alkaliphilic Cyanobacterium for Biomass Production with Direct Air CO2capture. J. CO2 Util. 2023, 69 (October 2022), 102399. 10.1016/j.jcou.2023.102399.

(24) Paerl, H. W.; Otten, T. G. Harmful Cyanobacterial Blooms: Causes, Consequences, and Controls. Microb. Ecol. 2013, 65 (4), 995–1010. 10.1007/s00248-012-0159-y.

(25) Ma, M.; Wei, C.; Huang, W.; He, Y.; Gong, Y.; Hu, Q. A Systematic Review of the Predatory Contaminant Poterioochromonas in Microalgal Culture. J. Appl. Phycol. 2023, 35 (3), 1103–1114. 10.1007/s10811-023-02941-0.

(26) Kilham, S. S.; Kreeger, D. A.; Lynn, S. G.; Goulden, C. E.; Herrera, L. COMBO: A Defined Freshwater Culture Medium for Algae and Zooplankton. Hydrobiologia 1998, 377 (1–3), 147–159. 10.1023/a:1003231628456.

(27) R Core Team. R: A Language and Environment for Statistical Computing. Vienna, Austria 2020. https://www.r-project.org/.

(28) Petzoldt, T. Estimation of Growth Rates with Package Growthrates. 2019.

(29) Kremer, C. T. GrowthTools. Zenodo February 2020. 10.5281/zenodo.3634918.

(30) Gardner, R. D.; Lohman, E.; Gerlach, R.; Cooksey, K. E.; Peyton, B. M. Comparison of CO2 and Bicarbonate as Inorganic Carbon Sources for Triacylglycerol and Starch Accumulation in Chlamydomonas Reinhardtii. Biotechnol. Bioeng. 2013, 110 (1), 87–96. 10.1002/bit.24592.

(31) Gardner, R. D.; Cooksey, K. E.; Mus, F.; Macur, R.; Moll, K.; Eustance, E.; Carlson, R. P.; Gerlach, R.; Fields, M. W.; Peyton, B. M. Use of Sodium Bicarbonate to Stimulate Triacylglycerol Accumulation in the Chlorophyte Scenedesmus Sp. and the Diatom Phaeodactylum Tricornutum. J. Appl. Phycol. 2012, 24 (5), 1311–1320. 10.1007/s10811-011-9782-0.

(32) Nunez, M.; Quigg, A. Changes in Growth and Composition of the Marine Microalgae Phaeodactylum Tricornutum and Nannochloropsis Salina in Response to Changing Sodium Bicarbonate Concentrations. J. Appl. Phycol. 2016, 28 (4), 2123–2138. 10.1007/s10811-015-0746-7.

