## Supplementary tables and figures for "Hunting for extremophiles: a systematic screening of freshwater microalgae for tolerance to high pH and high alkalinity cultivation"

ORCID IDs - PT: 0000-0002-7259-5766; RG: 0000-0002-7669-3072; AN: 0000-0003-4561-0163

Contents:

**A. Supplementary Tables**

**B. Supplementary Figures**

### Supplementary Tables

Table S1. Post hoc Games Howell test results showing differences in maximum fluorescence between treatments in Phase 1. “estimate” refers to the estimated mean difference of the first treatment versus the second treatment listed in the “contrast” column (i.e., the negative estimate in the first row means the pH & alkalinity treatment had lower max. fluorescence than the control). “conf.low” and “conf.high” show bounds of 95% confidence intervals; “p.adj” shows adjusted p values, and “p.adj.signif” shows the significance level of the contrast (ns: p>0.05, *: p<0.05; **p<0.01, ***p<0.001, ****p<0.0001 ).

| strain | contrast | estimate | conf.low | conf.high | p.adj | p.adj.signif |
| --- | --- | --- | --- | --- | --- | --- |
| Anabaena planctonica (NIVA-CYA 651) | pH & Alkalinity vs Control | -0.0762 | -0.1523 | -2.00E-04 | 0.05 | * |
| Anabaena planctonica (NIVA-CYA 651) | Salinity vs Control | 0.8208 | 0.7444 | 0.8973 | 0 | **** |
| Anabaena planctonica (NIVA-CYA 651) | Salinity vs pH & Alkalinity | 0.8971 | 0.8323 | 0.9618 | 0 | **** |
| Ankistrodesmus falcatus (SAG 202-2) | pH & Alkalinity vs Control | 0.2376 | 0.121 | 0.3542 | 0.002 | ** |
| Ankistrodesmus falcatus (SAG 202-2) | Salinity vs Control | -0.2011 | -0.3195 | -0.0827 | 0.01 | ** |
| Ankistrodesmus falcatus (SAG 202-2) | Salinity vs pH & Alkalinity | -0.4387 | -0.5218 | -0.3556 | 1.00E-04 | *** |
| Aphanizomenon flos-aquae (NIVA-CYA 681) | pH & Alkalinity vs Control | 0.0026 | -0.0394 | 0.0446 | 0.979 | ns |
| Aphanizomenon flos-aquae (NIVA-CYA 681) | Salinity vs Control | 0.0044 | -0.0421 | 0.0509 | 0.955 | ns |
| Aphanizomenon flos-aquae (NIVA-CYA 681) | Salinity vs pH & Alkalinity | 0.0017 | -0.0408 | 0.0443 | 0.991 | ns |
| Botryococcus braunii (CCAP 807/1) | pH & Alkalinity vs Control | 0.1642 | 0.0663 | 0.262 | 0.005 | ** |
| Botryococcus braunii (CCAP 807/1) | Salinity vs Control | 0.0046 | -0.2104 | 0.2195 | 0.997 | ns |
| Botryococcus braunii (CCAP 807/1) | Salinity vs pH & Alkalinity | -0.1596 | -0.3744 | 0.0552 | 0.117 | ns |
| Botryococcus braunii (CCAP 807/2) | pH & Alkalinity vs Control | -0.0791 | -0.1851 | 0.0269 | 0.106 | ns |
| Botryococcus braunii (CCAP 807/2) | Salinity vs Control | -0.1175 | -0.2083 | -0.0266 | 0.024 | * |
| Botryococcus braunii (CCAP 807/2) | Salinity vs pH & Alkalinity | -0.0384 | -0.1444 | 0.0675 | 0.539 | ns |
| Botryococcus braunii (EAWAG isolate PT1) | pH & Alkalinity vs Control | -0.4223 | -0.6865 | -0.1581 | 0.008 | ** |
| Botryococcus braunii (EAWAG isolate PT1) | Salinity vs Control | -0.3066 | -0.4748 | -0.1385 | 0.004 | ** |
| Botryococcus braunii (EAWAG isolate PT1) | Salinity vs pH & Alkalinity | 0.1157 | -0.1494 | 0.3808 | 0.366 | ns |
| Botryococcus braunii (NIES 2199) | pH & Alkalinity vs Control | 0.0568 | -0.0106 | 0.1241 | 0.09 | ns |
| Botryococcus braunii (NIES 2199) | Salinity vs Control | -0.0655 | -0.1195 | -0.0114 | 0.023 | * |
| Botryococcus braunii (NIES 2199) | Salinity vs pH & Alkalinity | -0.1222 | -0.1876 | -0.0568 | 0.004 | ** |
| Botryococcus braunii (SAG 30.81) | pH & Alkalinity vs Control | 0.018 | -0.0686 | 0.1046 | 0.796 | ns |
| Botryococcus braunii (SAG 30.81) | Salinity vs Control | -0.1787 | -0.2693 | -0.0881 | 0.002 | ** |
| Botryococcus braunii (SAG 30.81) | Salinity vs pH & Alkalinity | -0.1967 | -0.2735 | -0.1199 | 6.00E-04 | *** |
| Botryococcus braunii (Showa) | pH & Alkalinity vs Control | -0.7966 | -1.0886 | -0.5046 | 4.00E-04 | *** |
| Botryococcus braunii (Showa) | Salinity vs Control | -0.1987 | -0.4547 | 0.0573 | 0.107 | ns |
| Botryococcus braunii (Showa) | Salinity vs pH & Alkalinity | 0.5979 | 0.3389 | 0.8569 | 0.002 | ** |
| Chlamydomonas reinhardtii (CC 1690) | pH & Alkalinity vs Control | -0.0314 | -0.2042 | 0.1414 | 0.767 | ns |
| Chlamydomonas reinhardtii (CC 1690) | Salinity vs Control | -0.0981 | -0.2699 | 0.0738 | 0.196 | ns |
| Chlamydomonas reinhardtii (CC 1690) | Salinity vs pH & Alkalinity | -0.0667 | -0.1056 | -0.0277 | 0.005 | ** |
| Chlamydomonas reinhardtii (CC 3060) | pH & Alkalinity vs Control | 0.7855 | 0.5522 | 1.0188 | 0.001 | *** |
| Chlamydomonas reinhardtii (CC 3060) | Salinity vs Control | -0.0057 | -0.2408 | 0.2294 | 0.995 | ns |
| Chlamydomonas reinhardtii (CC 3060) | Salinity vs pH & Alkalinity | -0.7912 | -0.8468 | -0.7357 | 0 | **** |
| Chlorella vulgaris (SAG 211-11b) | pH & Alkalinity vs Control | 0.3821 | 0.1924 | 0.5718 | 0.007 | ** |
| Chlorella vulgaris (SAG 211-11b) | Salinity vs Control | -0.1143 | -0.1333 | -0.0952 | 0 | **** |
| Chlorella vulgaris (SAG 211-11b) | Salinity vs pH & Alkalinity | -0.4964 | -0.6869 | -0.3058 | 0.003 | ** |
| Cosmarium botrytis (SAG 136.8) | pH & Alkalinity vs Control | 0.1384 | -0.3311 | 0.608 | 0.657 | ns |
| Cosmarium botrytis (SAG 136.8) | Salinity vs Control | -0.2452 | -0.6568 | 0.1664 | 0.212 | ns |
| Cosmarium botrytis (SAG 136.8) | Salinity vs pH & Alkalinity | -0.3836 | -0.7973 | 0.0301 | 0.064 | ns |
| Cryptomonas sp. (NIVA 3/09) | pH & Alkalinity vs Control | -0.4894 | -0.6057 | -0.3731 | 0 | **** |
| Cryptomonas sp. (NIVA 3/09) | Salinity vs Control | -0.4106 | -0.8361 | 0.015 | 0.055 | ns |
| Cryptomonas sp. (NIVA 3/09) | Salinity vs pH & Alkalinity | 0.0789 | -0.3528 | 0.5105 | 0.768 | ns |
| Cyclotella meneghiniana (SAG 1020-1a) | pH & Alkalinity vs Control | 1.0163 | 0.9194 | 1.1133 | 0 | **** |
| Cyclotella meneghiniana (SAG 1020-1a) | Salinity vs Control | 0.6915 | 0.58 | 0.803 | 0 | **** |
| Cyclotella meneghiniana (SAG 1020-1a) | Salinity vs pH & Alkalinity | -0.3249 | -0.4237 | -0.226 | 4.00E-04 | *** |
| Cystodinium sp. (SAG 59.87) | pH & Alkalinity vs Control | -0.87 | -1.1162 | -0.6238 | 5.00E-04 | *** |
| Cystodinium sp. (SAG 59.87) | Salinity vs Control | -0.4196 | -0.8534 | 0.0142 | 0.056 | ns |
| Cystodinium sp. (SAG 59.87) | Salinity vs pH & Alkalinity | 0.4504 | -0.0076 | 0.9084 | 0.052 | ns |
| Desmodesmus abundans (EAWAG isolate GM26) | pH & Alkalinity vs Control | -0.1968 | -0.2693 | -0.1244 | 0.001 | *** |
| Desmodesmus abundans (EAWAG isolate GM26) | Salinity vs Control | -0.3423 | -0.4155 | -0.269 | 0 | **** |
| Desmodesmus abundans (EAWAG isolate GM26) | Salinity vs pH & Alkalinity | -0.1454 | -0.1959 | -0.095 | 4.00E-04 | *** |
| Desmodesmus armatus (EAWAG isolate L t0) | pH & Alkalinity vs Control | 0.3685 | 0.3301 | 0.407 | 0 | **** |
| Desmodesmus armatus (EAWAG isolate L t0) | Salinity vs Control | -0.2052 | -0.249 | -0.1614 | 0 | **** |
| Desmodesmus armatus (EAWAG isolate L t0) | Salinity vs pH & Alkalinity | -0.5738 | -0.6182 | -0.5293 | 0 | **** |
| Fistulifera saprophila (EAWAG isolate GM16) | pH & Alkalinity vs Control | 1.3991 | 0.0818 | 2.7163 | 0.042 | * |
| Fistulifera saprophila (EAWAG isolate GM16) | Salinity vs Control | 0.0632 | -1.2131 | 1.3395 | 0.983 | ns |
| Fistulifera saprophila (EAWAG isolate GM16) | Salinity vs pH & Alkalinity | -1.3358 | -1.8805 | -0.7912 | 0.001 | *** |
| Fragilaria capucina (CCAC 2678B) | pH & Alkalinity vs Control | -0.0793 | -0.3438 | 0.1852 | 0.557 | ns |
| Fragilaria capucina (CCAC 2678B) | Salinity vs Control | -0.0691 | -0.329 | 0.1908 | 0.692 | ns |
| Fragilaria capucina (CCAC 2678B) | Salinity vs pH & Alkalinity | 0.0102 | -0.1656 | 0.186 | 0.978 | ns |
| Fragilaria crotonensis (EAWAG isolate GM2) | pH & Alkalinity vs Control | -0.042 | -0.8779 | 0.7938 | 0.977 | ns |
| Fragilaria crotonensis (EAWAG isolate GM2) | Salinity vs Control | -0.1313 | -0.9689 | 0.7064 | 0.81 | ns |
| Fragilaria crotonensis (EAWAG isolate GM2) | Salinity vs pH & Alkalinity | -0.0892 | -0.2126 | 0.0342 | 0.146 | ns |
| Fragilaria mesolepta (EAWAG isolate GM15) | pH & Alkalinity vs Control | -0.8985 | -1.0699 | -0.727 | 0 | **** |
| Fragilaria mesolepta (EAWAG isolate GM15) | Salinity vs Control | -0.9421 | -1.0981 | -0.7862 | 1.00E-04 | **** |
| Fragilaria mesolepta (EAWAG isolate GM15) | Salinity vs pH & Alkalinity | -0.0436 | -0.1913 | 0.104 | 0.594 | ns |
| Kirchneriella subcapitata (SAG 12.81) | pH & Alkalinity vs Control | 0.4008 | 0.3304 | 0.4713 | 1.00E-04 | *** |
| Kirchneriella subcapitata (SAG 12.81) | Salinity vs Control | -0.1553 | -0.1881 | -0.1225 | 0 | **** |
| Kirchneriella subcapitata (SAG 12.81) | Salinity vs pH & Alkalinity | -0.5561 | -0.6251 | -0.4872 | 0 | **** |
| Lagerheimia hindakii (SAG 11.92) | pH & Alkalinity vs Control | 0.1732 | 0.1254 | 0.221 | 1.00E-04 | *** |
| Lagerheimia hindakii (SAG 11.92) | Salinity vs Control | -0.1808 | -0.4989 | 0.1374 | 0.19 | ns |
| Lagerheimia hindakii (SAG 11.92) | Salinity vs pH & Alkalinity | -0.354 | -0.6697 | -0.0383 | 0.037 | * |
| Lagerheimia subsalsa (EAWAG isolate GM37) | pH & Alkalinity vs Control | 0.4609 | 0.0437 | 0.8782 | 0.038 | * |
| Lagerheimia subsalsa (EAWAG isolate GM37) | Salinity vs Control | -0.5407 | -0.9463 | -0.1351 | 0.02 | * |
| Lagerheimia subsalsa (EAWAG isolate GM37) | Salinity vs pH & Alkalinity | -1.0016 | -1.1556 | -0.8477 | 0 | **** |
| Medakamo hakoo (NIES 4000) | pH & Alkalinity vs Control | 0.3372 | 0.2832 | 0.3913 | 0 | **** |
| Medakamo hakoo (NIES 4000) | Salinity vs Control | -0.0349 | -0.0892 | 0.0194 | 0.188 | ns |
| Medakamo hakoo (NIES 4000) | Salinity vs pH & Alkalinity | -0.3721 | -0.4078 | -0.3364 | 0 | **** |
| Messastrum gracile (EAWAG isolate GM45) | pH & Alkalinity vs Control | -0.0035 | -0.1667 | 0.1596 | 0.997 | ns |
| Messastrum gracile (EAWAG isolate GM45) | Salinity vs Control | -0.0122 | -0.1796 | 0.1552 | 0.961 | ns |
| Messastrum gracile (EAWAG isolate GM45) | Salinity vs pH & Alkalinity | -0.0087 | -0.0883 | 0.0709 | 0.937 | ns |
| Micractinium pusillum (CCAP 231/1) | pH & Alkalinity vs Control | -0.0923 | -0.14 | -0.0446 | 0.006 | ** |
| Micractinium pusillum (CCAP 231/1) | Salinity vs Control | 0.0373 | -0.007 | 0.0815 | 0.081 | ns |
| Micractinium pusillum (CCAP 231/1) | Salinity vs pH & Alkalinity | 0.1296 | 0.0787 | 0.1805 | 6.00E-04 | *** |
| Microcystis aeruginosa (EAWAG isolate Greifensee) | pH & Alkalinity vs Control | -0.9326 | -1.0146 | -0.8507 | 0 | **** |
| Microcystis aeruginosa (EAWAG isolate Greifensee) | Salinity vs Control | -0.7785 | -0.8166 | -0.7403 | 0 | **** |
| Microcystis aeruginosa (EAWAG isolate Greifensee) | Salinity vs pH & Alkalinity | 0.1542 | 0.077 | 0.2313 | 0.004 | ** |
| Microcystis aeruginosa (PCC 7806) | pH & Alkalinity vs Control | 0.3855 | 0.2948 | 0.4762 | 1.00E-04 | **** |
| Microcystis aeruginosa (PCC 7806) | Salinity vs Control | 0.3133 | 0.03 | 0.5967 | 0.038 | * |
| Microcystis aeruginosa (PCC 7806) | Salinity vs pH & Alkalinity | -0.0722 | -0.3488 | 0.2045 | 0.633 | ns |
| Nannochloropsis limnetica (SAG 18.99) | pH & Alkalinity vs Control | 0.1276 | 0.0195 | 0.2357 | 0.031 | * |
| Nannochloropsis limnetica (SAG 18.99) | Salinity vs Control | -0.1232 | -0.1599 | -0.0865 | 1.00E-04 | *** |
| Nannochloropsis limnetica (SAG 18.99) | Salinity vs pH & Alkalinity | -0.2508 | -0.3575 | -0.144 | 0.003 | ** |
| Nannochloropsis oculata (SAG 38.85) | pH & Alkalinity vs Control | 0.6719 | 0.5984 | 0.7454 | 0 | **** |
| Nannochloropsis oculata (SAG 38.85) | Salinity vs Control | 0.7178 | 0.6461 | 0.7896 | 0 | **** |
| Nannochloropsis oculata (SAG 38.85) | Salinity vs pH & Alkalinity | 0.0459 | -0.0364 | 0.1282 | 0.276 | ns |
| Oocystella heteromucosa (EAWAG isolate GM35) | pH & Alkalinity vs Control | 0.2114 | -0.1338 | 0.5566 | 0.165 | ns |
| Oocystella heteromucosa (EAWAG isolate GM35) | Salinity vs Control | -0.474 | -0.734 | -0.2139 | 0.009 | ** |
| Oocystella heteromucosa (EAWAG isolate GM35) | Salinity vs pH & Alkalinity | -0.6854 | -1.0164 | -0.3544 | 0.002 | ** |
| Oocystis solitaria (SAG 83.8) | pH & Alkalinity vs Control | 0.101 | -0.0522 | 0.2543 | 0.183 | ns |
| Oocystis solitaria (SAG 83.8) | Salinity vs Control | -0.0115 | -0.1461 | 0.1231 | 0.963 | ns |
| Oocystis solitaria (SAG 83.8) | Salinity vs pH & Alkalinity | -0.1125 | -0.2696 | 0.0445 | 0.148 | ns |
| Oocystis sp. (EAWAG isolate L t0) | pH & Alkalinity vs Control | 0.3866 | 0.3017 | 0.4714 | 0 | **** |
| Oocystis sp. (EAWAG isolate L t0) | Salinity vs Control | -0.1716 | -0.2556 | -0.0876 | 0.002 | ** |
| Oocystis sp. (EAWAG isolate L t0) | Salinity vs pH & Alkalinity | -0.5581 | -0.6369 | -0.4794 | 0 | **** |
| Pandorina unicocca (EAWAG isolate GM17) | pH & Alkalinity vs Control | -1.115 | -1.2154 | -1.0145 | 0 | **** |
| Pandorina unicocca (EAWAG isolate GM17) | Salinity vs Control | -1.2109 | -1.354 | -1.0678 | 0 | **** |
| Pandorina unicocca (EAWAG isolate GM17) | Salinity vs pH & Alkalinity | -0.0959 | -0.2398 | 0.0479 | 0.171 | ns |
| Pediastrum boryanum (EAWAG isolate GM20) | pH & Alkalinity vs Control | 0.833 | 0.4135 | 1.2525 | 0.006 | ** |
| Pediastrum boryanum (EAWAG isolate GM20) | Salinity vs Control | 0.3672 | 0.2661 | 0.4683 | 1.00E-04 | *** |
| Pediastrum boryanum (EAWAG isolate GM20) | Salinity vs pH & Alkalinity | -0.4658 | -0.8911 | -0.0405 | 0.039 | * |
| Pediastrum boryanum (SAG 87.81) | pH & Alkalinity vs Control | -0.2798 | -0.426 | -0.1337 | 0.005 | ** |
| Pediastrum boryanum (SAG 87.81) | Salinity vs Control | -0.2473 | -0.3942 | -0.1005 | 0.009 | ** |
| Pediastrum boryanum (SAG 87.81) | Salinity vs pH & Alkalinity | 0.0325 | -0.0312 | 0.0962 | 0.329 | ns |
| Pediastrum duplex (EAWAG isolate GM22) | pH & Alkalinity vs Control | -0.7912 | -0.9068 | -0.6756 | 0 | **** |
| Pediastrum duplex (EAWAG isolate GM22) | Salinity vs Control | -0.7798 | -0.9742 | -0.5853 | 1.00E-04 | *** |
| Pediastrum duplex (EAWAG isolate GM22) | Salinity vs pH & Alkalinity | 0.0114 | -0.1857 | 0.2085 | 0.977 | ns |
| Pediastrum duplex (SAG 261-2) | pH & Alkalinity vs Control | 0.5113 | 0.1585 | 0.864 | 0.018 | * |
| Pediastrum duplex (SAG 261-2) | Salinity vs Control | -0.0759 | -0.2269 | 0.0751 | 0.246 | ns |
| Pediastrum duplex (SAG 261-2) | Salinity vs pH & Alkalinity | -0.5872 | -0.9131 | -0.2613 | 0.006 | ** |
| Planktothrix rubescens (NIVA-CYA 619) | pH & Alkalinity vs Control | 0.3342 | -0.0838 | 0.7522 | 0.088 | ns |
| Planktothrix rubescens (NIVA-CYA 619) | Salinity vs Control | 0.0477 | -0.3726 | 0.4679 | 0.889 | ns |
| Planktothrix rubescens (NIVA-CYA 619) | Salinity vs pH & Alkalinity | -0.2865 | -0.324 | -0.249 | 0 | **** |
| Poterioochromonas malhamensis (CCAP 933/1C) | pH & Alkalinity vs Control | 0.5737 | -0.0141 | 1.1615 | 0.055 | ns |
| Poterioochromonas malhamensis (CCAP 933/1C) | Salinity vs Control | 0.487 | -0.0325 | 1.0065 | 0.06 | ns |
| Poterioochromonas malhamensis (CCAP 933/1C) | Salinity vs pH & Alkalinity | -0.0867 | -0.6456 | 0.4722 | 0.829 | ns |
| Scenedesmus acuminatus (SAG 38.81) | pH & Alkalinity vs Control | -0.0243 | -0.0864 | 0.0379 | 0.474 | ns |
| Scenedesmus acuminatus (SAG 38.81) | Salinity vs Control | -0.1563 | -0.247 | -0.0655 | 0.007 | ** |
| Scenedesmus acuminatus (SAG 38.81) | Salinity vs pH & Alkalinity | -0.132 | -0.2234 | -0.0406 | 0.012 | * |
| Scenedesmus armatus (EAWAG isolate GM28) | pH & Alkalinity vs Control | 0.3769 | 0.2557 | 0.4981 | 5.00E-04 | *** |
| Scenedesmus armatus (EAWAG isolate GM28) | Salinity vs Control | -0.2465 | -0.3164 | -0.1765 | 1.00E-04 | *** |
| Scenedesmus armatus (EAWAG isolate GM28) | Salinity vs pH & Alkalinity | -0.6234 | -0.7452 | -0.5015 | 1.00E-04 | **** |
| Scenedesmus ellipticus (EAWAG isolate GM25) | pH & Alkalinity vs Control | 0.2926 | 0.2155 | 0.3697 | 5.00E-04 | *** |
| Scenedesmus ellipticus (EAWAG isolate GM25) | Salinity vs Control | -0.3619 | -0.442 | -0.2818 | 0 | **** |
| Scenedesmus ellipticus (EAWAG isolate GM25) | Salinity vs pH & Alkalinity | -0.6545 | -0.7194 | -0.5896 | 0 | **** |
| Staurastrum punctulatum (SAG 679-1) | pH & Alkalinity vs Control | -0.7828 | -0.9325 | -0.6332 | 0 | **** |
| Staurastrum punctulatum (SAG 679-1) | Salinity vs Control | -0.7295 | -1.0042 | -0.4547 | 0.001 | *** |
| Staurastrum punctulatum (SAG 679-1) | Salinity vs pH & Alkalinity | 0.0534 | -0.227 | 0.3337 | 0.78 | ns |
| Synechococcus elongatus (EAWAG isolate GM48) | pH & Alkalinity vs Control | 0.1115 | 0.0629 | 0.1601 | 0.004 | ** |
| Synechococcus elongatus (EAWAG isolate GM48) | Salinity vs Control | 0.2635 | 0.2146 | 0.3124 | 0 | **** |
| Synechococcus elongatus (EAWAG isolate GM48) | Salinity vs pH & Alkalinity | 0.152 | 0.1116 | 0.1924 | 7.00E-04 | *** |
| Synechocystis sp. (PCC 6803) | pH & Alkalinity vs Control | 0.2459 | 0.003 | 0.4889 | 0.048 | * |
| Synechocystis sp. (PCC 6803) | Salinity vs Control | 0.0222 | -0.2115 | 0.2559 | 0.93 | ns |
| Synechocystis sp. (PCC 6803) | Salinity vs pH & Alkalinity | -0.2237 | -0.2836 | -0.1638 | 7.00E-04 | *** |
| Tetraedron minimum (SAG 44.81) | pH & Alkalinity vs Control | 0.025 | -0.2801 | 0.3301 | 0.94 | ns |
| Tetraedron minimum (SAG 44.81) | Salinity vs Control | -0.3152 | -0.6102 | -0.0203 | 0.041 | * |
| Tetraedron minimum (SAG 44.81) | Salinity vs pH & Alkalinity | -0.3402 | -0.4108 | -0.2696 | 2.00E-04 | *** |

Table S2. Post hoc Games Howell test results showing differences in maximum fluorescence between treatments in Phase 2. “estimate” refers to the estimated mean difference of the first treatment versus the second treatment listed in the “contrast” column (i.e., the negative estimate in the first row means the pH & alkalinity treatment had lower max. fluorescence than the control). “conf.low” and “conf.high” show bounds of 95% confidence intervals; “p.adj” shows adjusted p values, and “p.adj.signif” shows the significance level of the contrast (ns: p>0.05, *: p<0.05; **p<0.01, ***p<0.001, ****p<0.0001 ).

| strain | contrast | estimate | conf.low | conf.high | p.adj | p.adj.signif |
| --- | --- | --- | --- | --- | --- | --- |
| Anabaena planctonica (NIVA-CYA 651) | pH & Alkalinity vs Control | -4.3147 | -5.2263 | -3.4031 | 3.00E-04 | *** |
| Anabaena planctonica (NIVA-CYA 651) | Salinity vs Control | -2.9382 | -7.0454 | 1.1689 | 0.119 | ns |
| Anabaena planctonica (NIVA-CYA 651) | Salinity vs pH & Alkalinity | 1.3765 | -2.8612 | 5.6141 | 0.464 | ns |
| Ankistrodesmus falcatus (SAG 202-2) | pH & Alkalinity vs Control | -1.7618 | -2.0567 | -1.4669 | 1.00E-04 | **** |
| Ankistrodesmus falcatus (SAG 202-2) | Salinity vs Control | -0.7747 | -0.9161 | -0.6333 | 0 | **** |
| Ankistrodesmus falcatus (SAG 202-2) | Salinity vs pH & Alkalinity | 0.9871 | 0.6942 | 1.28 | 5.00E-04 | *** |
| Aphanizomenon flos-aquae (NIVA-CYA 681) | pH & Alkalinity vs Control | -4.7031 | -5.0156 | -4.3907 | 0 | **** |
| Aphanizomenon flos-aquae (NIVA-CYA 681) | Salinity vs Control | -1.6606 | -2.4125 | -0.9086 | 0.003 | ** |
| Aphanizomenon flos-aquae (NIVA-CYA 681) | Salinity vs pH & Alkalinity | 3.0426 | 2.2457 | 3.8395 | 9.00E-04 | *** |
| Botryococcus braunii (CCAP 807/1) | pH & Alkalinity vs Control | -1.8172 | -2.2813 | -1.3532 | 8.00E-04 | *** |
| Botryococcus braunii (CCAP 807/1) | Salinity vs Control | -0.0582 | -0.2659 | 0.1494 | 0.62 | ns |
| Botryococcus braunii (CCAP 807/1) | Salinity vs pH & Alkalinity | 1.759 | 1.3213 | 2.1967 | 2.00E-04 | *** |
| Botryococcus braunii (CCAP 807/2) | pH & Alkalinity vs Control | -1.253 | -1.4533 | -1.0527 | 0 | **** |
| Botryococcus braunii (CCAP 807/2) | Salinity vs Control | -0.2653 | -0.4324 | -0.0982 | 0.007 | ** |
| Botryococcus braunii (CCAP 807/2) | Salinity vs pH & Alkalinity | 0.9878 | 0.801 | 1.1745 | 0 | **** |
| Botryococcus braunii (EAWAG isolate PT1) | pH & Alkalinity vs Control | -2.9165 | -3.3135 | -2.5195 | 0 | **** |
| Botryococcus braunii (EAWAG isolate PT1) | Salinity vs Control | -0.9922 | -1.2786 | -0.7057 | 1.00E-04 | *** |
| Botryococcus braunii (EAWAG isolate PT1) | Salinity vs pH & Alkalinity | 1.9243 | 1.5217 | 2.3269 | 0 | **** |
| Botryococcus braunii (NIES 2199) | pH & Alkalinity vs Control | -2.2304 | -2.3376 | -2.1232 | 0 | **** |
| Botryococcus braunii (NIES 2199) | Salinity vs Control | -0.4489 | -0.574 | -0.3237 | 1.00E-04 | *** |
| Botryococcus braunii (NIES 2199) | Salinity vs pH & Alkalinity | 1.7816 | 1.6577 | 1.9054 | 0 | **** |
| Botryococcus braunii (SAG 30.81) | pH & Alkalinity vs Control | -0.3984 | -0.6909 | -0.1058 | 0.017 | * |
| Botryococcus braunii (SAG 30.81) | Salinity vs Control | -0.1318 | -0.3203 | 0.0567 | 0.157 | ns |
| Botryococcus braunii (SAG 30.81) | Salinity vs pH & Alkalinity | 0.2666 | -0.0262 | 0.5594 | 0.069 | ns |
| Botryococcus braunii (Showa) | pH & Alkalinity vs Control | -3.4367 | -4.1691 | -2.7043 | 5.00E-04 | *** |
| Botryococcus braunii (Showa) | Salinity vs Control | -1.6073 | -2.29 | -0.9246 | 0.002 | ** |
| Botryococcus braunii (Showa) | Salinity vs pH & Alkalinity | 1.8294 | 1.4218 | 2.237 | 4.00E-04 | *** |
| Chlamydomonas reinhardtii (CC 1690) | pH & Alkalinity vs Control | -2.8908 | -3.0969 | -2.6846 | 0 | **** |
| Chlamydomonas reinhardtii (CC 1690) | Salinity vs Control | -0.8191 | -0.9043 | -0.7338 | 0 | **** |
| Chlamydomonas reinhardtii (CC 1690) | Salinity vs pH & Alkalinity | 2.0717 | 1.8641 | 2.2793 | 0 | **** |
| Chlamydomonas reinhardtii (CC 3060) | pH & Alkalinity vs Control | -4.4377 | -6.0459 | -2.8295 | 0.003 | ** |
| Chlamydomonas reinhardtii (CC 3060) | Salinity vs Control | -0.6351 | -0.9136 | -0.3565 | 0.001 | *** |
| Chlamydomonas reinhardtii (CC 3060) | Salinity vs pH & Alkalinity | 3.8026 | 2.2125 | 5.3927 | 0.004 | ** |
| Chlorella vulgaris (SAG 211-11b) | pH & Alkalinity vs Control | -0.9304 | -1.3047 | -0.5561 | 0.004 | ** |
| Chlorella vulgaris (SAG 211-11b) | Salinity vs Control | 0.0094 | -0.1157 | 0.1346 | 0.952 | ns |
| Chlorella vulgaris (SAG 211-11b) | Salinity vs pH & Alkalinity | 0.9399 | 0.5879 | 1.2918 | 0.002 | ** |
| Cosmarium botrytis (SAG 136.8) | pH & Alkalinity vs Control | -4.0541 | -4.7918 | -3.3164 | 1.00E-04 | **** |
| Cosmarium botrytis (SAG 136.8) | Salinity vs Control | -2.5819 | -2.9894 | -2.1744 | 0 | **** |
| Cosmarium botrytis (SAG 136.8) | Salinity vs pH & Alkalinity | 1.4721 | 0.7335 | 2.2108 | 0.004 | ** |
| Cryptomonas sp. (NIVA 3/09) | pH & Alkalinity vs Control | -2.5528 | -4.785 | -0.3206 | 0.035 | * |
| Cryptomonas sp. (NIVA 3/09) | Salinity vs Control | -2.2383 | -4.3596 | -0.117 | 0.042 | * |
| Cryptomonas sp. (NIVA 3/09) | Salinity vs pH & Alkalinity | 0.3145 | -0.499 | 1.1281 | 0.445 | ns |
| Cyclotella meneghiniana (SAG 1020-1a) | pH & Alkalinity vs Control | -2.5442 | -2.8304 | -2.258 | 0 | **** |
| Cyclotella meneghiniana (SAG 1020-1a) | Salinity vs Control | -0.0247 | -0.2963 | 0.2469 | 0.938 | ns |
| Cyclotella meneghiniana (SAG 1020-1a) | Salinity vs pH & Alkalinity | 2.5195 | 2.4338 | 2.6052 | 0 | **** |
| Cystodinium sp. (SAG 59.87) | pH & Alkalinity vs Control | -1.6612 | -2.2966 | -1.0258 | 0.002 | ** |
| Cystodinium sp. (SAG 59.87) | Salinity vs Control | -0.8278 | -1.3722 | -0.2834 | 0.012 | * |
| Cystodinium sp. (SAG 59.87) | Salinity vs pH & Alkalinity | 0.8334 | 0.1629 | 1.5039 | 0.021 | * |
| Desmodesmus abundans (EAWAG isolate GM26) | pH & Alkalinity vs Control | -0.1615 | -0.3149 | -0.0082 | 0.042 | * |
| Desmodesmus abundans (EAWAG isolate GM26) | Salinity vs Control | -0.2109 | -0.3019 | -0.1198 | 0.001 | *** |
| Desmodesmus abundans (EAWAG isolate GM26) | Salinity vs pH & Alkalinity | -0.0493 | -0.2026 | 0.1039 | 0.565 | ns |
| Desmodesmus armatus (EAWAG isolate L t0) | pH & Alkalinity vs Control | -0.6657 | -0.7379 | -0.5934 | 0 | **** |
| Desmodesmus armatus (EAWAG isolate L t0) | Salinity vs Control | -0.2871 | -0.3126 | -0.2616 | 0 | **** |
| Desmodesmus armatus (EAWAG isolate L t0) | Salinity vs pH & Alkalinity | 0.3786 | 0.3087 | 0.4484 | 1.00E-04 | *** |
| Fistulifera saprophila (EAWAG isolate GM16) | pH & Alkalinity vs Control | -4.6057 | -8.0122 | -1.1991 | 0.022 | * |
| Fistulifera saprophila (EAWAG isolate GM16) | Salinity vs Control | -0.763 | -2.8318 | 1.3058 | 0.395 | ns |
| Fistulifera saprophila (EAWAG isolate GM16) | Salinity vs pH & Alkalinity | 3.8427 | 0.7208 | 6.9647 | 0.023 | * |
| Fragilaria capucina (CCAC 2678B) | pH & Alkalinity vs Control | -3.0175 | -6.5177 | 0.4828 | 0.073 | ns |
| Fragilaria capucina (CCAC 2678B) | Salinity vs Control | -3.1149 | -6.609 | 0.3792 | 0.067 | ns |
| Fragilaria capucina (CCAC 2678B) | Salinity vs pH & Alkalinity | -0.0974 | -0.3873 | 0.1924 | 0.581 | ns |
| Fragilaria crotonensis (EAWAG isolate GM2) | pH & Alkalinity vs Control | -2.4309 | -2.8544 | -2.0074 | 1.00E-04 | **** |
| Fragilaria crotonensis (EAWAG isolate GM2) | Salinity vs Control | -2.3659 | -2.5532 | -2.1786 | 0 | **** |
| Fragilaria crotonensis (EAWAG isolate GM2) | Salinity vs pH & Alkalinity | 0.065 | -0.3667 | 0.4967 | 0.848 | ns |
| Fragilaria mesolepta (EAWAG isolate GM15) | pH & Alkalinity vs Control | -4.2354 | -4.5297 | -3.9411 | 0 | **** |
| Fragilaria mesolepta (EAWAG isolate GM15) | Salinity vs Control | -4.4203 | -4.7665 | -4.0742 | 0 | **** |
| Fragilaria mesolepta (EAWAG isolate GM15) | Salinity vs pH & Alkalinity | -0.1849 | -0.5406 | 0.1707 | 0.315 | ns |
| Kirchneriella subcapitata (SAG 12.81) | pH & Alkalinity vs Control | -1.3731 | -1.439 | -1.3071 | 0 | **** |
| Kirchneriella subcapitata (SAG 12.81) | Salinity vs Control | -0.1572 | -0.2175 | -0.0969 | 0.001 | *** |
| Kirchneriella subcapitata (SAG 12.81) | Salinity vs pH & Alkalinity | 1.2159 | 1.1438 | 1.2879 | 0 | **** |
| Lagerheimia hindakii (SAG 11.92) | pH & Alkalinity vs Control | -1.4273 | -1.5084 | -1.3461 | 0 | **** |
| Lagerheimia hindakii (SAG 11.92) | Salinity vs Control | -0.2461 | -0.3513 | -0.1409 | 0.001 | *** |
| Lagerheimia hindakii (SAG 11.92) | Salinity vs pH & Alkalinity | 1.1811 | 1.0787 | 1.2836 | 0 | **** |
| Lagerheimia subsalsa (EAWAG isolate GM37) | pH & Alkalinity vs Control | -1.2853 | -1.7033 | -0.8673 | 0.002 | ** |
| Lagerheimia subsalsa (EAWAG isolate GM37) | Salinity vs Control | -0.7271 | -1.1189 | -0.3352 | 0.006 | ** |
| Lagerheimia subsalsa (EAWAG isolate GM37) | Salinity vs pH & Alkalinity | 0.5583 | 0.3755 | 0.741 | 0.001 | *** |
| Medakamo hakoo (NIES 4000) | pH & Alkalinity vs Control | -1.2016 | -1.2855 | -1.1177 | 0 | **** |
| Medakamo hakoo (NIES 4000) | Salinity vs Control | -0.2274 | -0.2938 | -0.1611 | 1.00E-04 | *** |
| Medakamo hakoo (NIES 4000) | Salinity vs pH & Alkalinity | 0.9742 | 0.8939 | 1.0544 | 0 | **** |
| Messastrum gracile (EAWAG isolate GM45) | pH & Alkalinity vs Control | -1.8183 | -2.126 | -1.5106 | 1.00E-04 | **** |
| Messastrum gracile (EAWAG isolate GM45) | Salinity vs Control | 0.0713 | -0.0467 | 0.1892 | 0.204 | ns |
| Messastrum gracile (EAWAG isolate GM45) | Salinity vs pH & Alkalinity | 1.8895 | 1.5667 | 2.2124 | 2.00E-04 | *** |
| Micractinium pusillum (CCAP 231/1) | pH & Alkalinity vs Control | -2.5993 | -3.0135 | -2.1851 | 1.00E-04 | *** |
| Micractinium pusillum (CCAP 231/1) | Salinity vs Control | -0.1548 | -0.264 | -0.0455 | 0.012 | * |
| Micractinium pusillum (CCAP 231/1) | Salinity vs pH & Alkalinity | 2.4445 | 2.0221 | 2.8669 | 3.00E-04 | *** |
| Microcystis aeruginosa (EAWAG isolate Greifensee) | pH & Alkalinity vs Control | -4.1129 | -4.2129 | -4.013 | 0 | **** |
| Microcystis aeruginosa (EAWAG isolate Greifensee) | Salinity vs Control | -1.0917 | -1.277 | -0.9064 | 0 | **** |
| Microcystis aeruginosa (EAWAG isolate Greifensee) | Salinity vs pH & Alkalinity | 3.0212 | 2.8305 | 3.212 | 0 | **** |
| Microcystis aeruginosa (PCC 7806) | pH & Alkalinity vs Control | -2.9838 | -3.2555 | -2.7121 | 0 | **** |
| Microcystis aeruginosa (PCC 7806) | Salinity vs Control | -0.3679 | -0.4492 | -0.2866 | 0 | **** |
| Microcystis aeruginosa (PCC 7806) | Salinity vs pH & Alkalinity | 2.616 | 2.3461 | 2.8859 | 0 | **** |
| Nannochloropsis limnetica (SAG 18.99) | pH & Alkalinity vs Control | -2.5367 | -2.6534 | -2.42 | 0 | **** |
| Nannochloropsis limnetica (SAG 18.99) | Salinity vs Control | -1.3775 | -1.4774 | -1.2776 | 0 | **** |
| Nannochloropsis limnetica (SAG 18.99) | Salinity vs pH & Alkalinity | 1.1591 | 1.0526 | 1.2657 | 0 | **** |
| Nannochloropsis oculata (SAG 38.85) | pH & Alkalinity vs Control | -1.9518 | -2.0434 | -1.8603 | 0 | **** |
| Nannochloropsis oculata (SAG 38.85) | Salinity vs Control | 0.2296 | 0.1848 | 0.2745 | 2.00E-04 | *** |
| Nannochloropsis oculata (SAG 38.85) | Salinity vs pH & Alkalinity | 2.1814 | 2.0835 | 2.2794 | 0 | **** |
| Oocystella heteromucosa (EAWAG isolate GM35) | pH & Alkalinity vs Control | -2.6559 | -3.0435 | -2.2683 | 1.00E-04 | *** |
| Oocystella heteromucosa (EAWAG isolate GM35) | Salinity vs Control | -2.8654 | -3.19 | -2.5409 | 0 | **** |
| Oocystella heteromucosa (EAWAG isolate GM35) | Salinity vs pH & Alkalinity | -0.2096 | -0.5876 | 0.1685 | 0.277 | ns |
| Oocystis solitaria (SAG 83.8) | pH & Alkalinity vs Control | -2.7447 | -3.0771 | -2.4122 | 0 | **** |
| Oocystis solitaria (SAG 83.8) | Salinity vs Control | -1.5293 | -1.7442 | -1.3144 | 1.00E-04 | **** |
| Oocystis solitaria (SAG 83.8) | Salinity vs pH & Alkalinity | 1.2154 | 0.8956 | 1.5351 | 1.00E-04 | *** |
| Oocystis sp. (EAWAG isolate L t0) | pH & Alkalinity vs Control | -2.2561 | -2.555 | -1.9572 | 1.00E-04 | **** |
| Oocystis sp. (EAWAG isolate L t0) | Salinity vs Control | -0.2899 | -0.3465 | -0.2333 | 0 | **** |
| Oocystis sp. (EAWAG isolate L t0) | Salinity vs pH & Alkalinity | 1.9662 | 1.6661 | 2.2663 | 2.00E-04 | *** |
| Pandorina unicocca (EAWAG isolate GM17) | pH & Alkalinity vs Control | -3.918 | -4.0196 | -3.8164 | 0 | **** |
| Pandorina unicocca (EAWAG isolate GM17) | Salinity vs Control | -3.7789 | -4.1808 | -3.377 | 0 | **** |
| Pandorina unicocca (EAWAG isolate GM17) | Salinity vs pH & Alkalinity | 0.1391 | -0.2643 | 0.5424 | 0.454 | ns |
| Pediastrum boryanum (EAWAG isolate GM20) | pH & Alkalinity vs Control | -2.2767 | -3.2491 | -1.3042 | 0.002 | ** |
| Pediastrum boryanum (EAWAG isolate GM20) | Salinity vs Control | -1.2439 | -1.7234 | -0.7645 | 5.00E-04 | *** |
| Pediastrum boryanum (EAWAG isolate GM20) | Salinity vs pH & Alkalinity | 1.0328 | 0.0646 | 2.0009 | 0.041 | * |
| Pediastrum boryanum (SAG 87.81) | pH & Alkalinity vs Control | -1.8061 | -2.2107 | -1.4015 | 4.00E-04 | *** |
| Pediastrum boryanum (SAG 87.81) | Salinity vs Control | -0.4287 | -0.5532 | -0.3043 | 1.00E-04 | *** |
| Pediastrum boryanum (SAG 87.81) | Salinity vs pH & Alkalinity | 1.3773 | 0.9682 | 1.7865 | 0.001 | *** |
| Pediastrum duplex (EAWAG isolate GM22) | pH & Alkalinity vs Control | -4.3651 | -4.8209 | -3.9093 | 0 | **** |
| Pediastrum duplex (EAWAG isolate GM22) | Salinity vs Control | -2.943 | -3.3728 | -2.5132 | 0 | **** |
| Pediastrum duplex (EAWAG isolate GM22) | Salinity vs pH & Alkalinity | 1.4221 | 0.9526 | 1.8916 | 2.00E-04 | *** |
| Pediastrum duplex (SAG 261-2) | pH & Alkalinity vs Control | -1.3662 | -2.2735 | -0.4588 | 0.016 | * |
| Pediastrum duplex (SAG 261-2) | Salinity vs Control | -0.2262 | -0.3261 | -0.1264 | 0.003 | ** |
| Pediastrum duplex (SAG 261-2) | Salinity vs pH & Alkalinity | 1.1399 | 0.2406 | 2.0393 | 0.026 | * |
| Planktothrix rubescens (NIVA-CYA 619) | pH & Alkalinity vs Control | -1.3883 | -1.6417 | -1.1349 | 0 | **** |
| Planktothrix rubescens (NIVA-CYA 619) | Salinity vs Control | -0.4429 | -0.6871 | -0.1988 | 0.004 | ** |
| Planktothrix rubescens (NIVA-CYA 619) | Salinity vs pH & Alkalinity | 0.9454 | 0.6655 | 1.2252 | 1.00E-04 | *** |
| Poterioochromonas malhamensis (CCAP 933/1C) | pH & Alkalinity vs Control | -2.4024 | -2.9561 | -1.8487 | 2.00E-04 | *** |
| Poterioochromonas malhamensis (CCAP 933/1C) | Salinity vs Control | 0.6003 | 0.039 | 1.1616 | 0.041 | * |
| Poterioochromonas malhamensis (CCAP 933/1C) | Salinity vs pH & Alkalinity | 3.0027 | 2.7196 | 3.2858 | 0 | **** |
| Scenedesmus acuminatus (SAG 38.81) | pH & Alkalinity vs Control | -3.8373 | -6.0662 | -1.6085 | 0.011 | * |
| Scenedesmus acuminatus (SAG 38.81) | Salinity vs Control | -0.2998 | -0.3676 | -0.2319 | 2.00E-04 | *** |
| Scenedesmus acuminatus (SAG 38.81) | Salinity vs pH & Alkalinity | 3.5375 | 1.3071 | 5.768 | 0.014 | * |
| Scenedesmus armatus (EAWAG isolate GM28) | pH & Alkalinity vs Control | -0.366 | -0.4498 | -0.2822 | 3.00E-04 | *** |
| Scenedesmus armatus (EAWAG isolate GM28) | Salinity vs Control | -0.045 | -0.0786 | -0.0114 | 0.015 | * |
| Scenedesmus armatus (EAWAG isolate GM28) | Salinity vs pH & Alkalinity | 0.3211 | 0.2367 | 0.4054 | 5.00E-04 | *** |
| Scenedesmus ellipticus (EAWAG isolate GM25) | pH & Alkalinity vs Control | -0.6637 | -0.8198 | -0.5076 | 1.00E-04 | **** |
| Scenedesmus ellipticus (EAWAG isolate GM25) | Salinity vs Control | -0.4089 | -0.5892 | -0.2285 | 0.001 | *** |
| Scenedesmus ellipticus (EAWAG isolate GM25) | Salinity vs pH & Alkalinity | 0.2548 | 0.0908 | 0.4188 | 0.009 | ** |
| Staurastrum punctulatum (SAG 679-1) | pH & Alkalinity vs Control | -3.5085 | -3.8099 | -3.2071 | 0 | **** |
| Staurastrum punctulatum (SAG 679-1) | Salinity vs Control | -3.542 | -3.9212 | -3.1627 | 0 | **** |
| Staurastrum punctulatum (SAG 679-1) | Salinity vs pH & Alkalinity | -0.0335 | -0.4151 | 0.3481 | 0.96 | ns |
| Synechococcus elongatus (EAWAG isolate GM48) | pH & Alkalinity vs Control | -0.9615 | -1.0781 | -0.845 | 0 | **** |
| Synechococcus elongatus (EAWAG isolate GM48) | Salinity vs Control | 0.4462 | 0.3708 | 0.5216 | 2.00E-04 | *** |
| Synechococcus elongatus (EAWAG isolate GM48) | Salinity vs pH & Alkalinity | 1.4077 | 1.2976 | 1.5179 | 0 | **** |
| Synechocystis sp. (PCC 6803) | pH & Alkalinity vs Control | 0.1364 | 0.0261 | 0.2467 | 0.026 | * |
| Synechocystis sp. (PCC 6803) | Salinity vs Control | 0.2517 | 0.1914 | 0.312 | 1.00E-04 | **** |
| Synechocystis sp. (PCC 6803) | Salinity vs pH & Alkalinity | 0.1153 | 0.0081 | 0.2225 | 0.039 | * |
| Tetraedron minimum (SAG 44.81) | pH & Alkalinity vs Control | -1.6503 | -1.7548 | -1.5458 | 0 | **** |
| Tetraedron minimum (SAG 44.81) | Salinity vs Control | -0.6059 | -0.692 | -0.5198 | 0 | **** |
| Tetraedron minimum (SAG 44.81) | Salinity vs pH & Alkalinity | 1.0444 | 0.9344 | 1.1543 | 0 | **** |

### Supplementary Figures

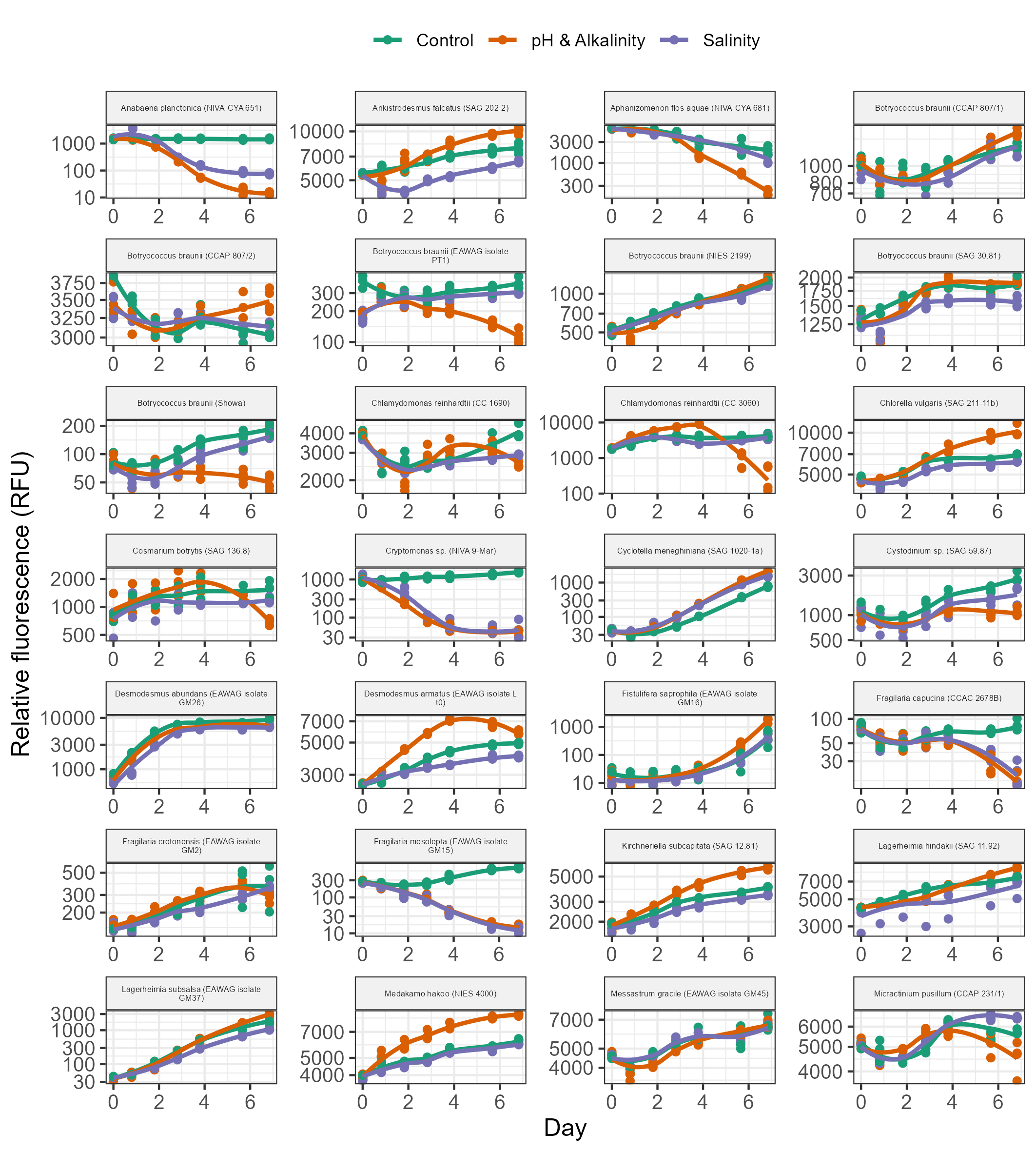

Fig. S1. Growth curves of all microalgae strains in Phase 1 (i.e., the acclimation phase to a mild increase in pH and alkalinity – pH 8.5 and 25 mM of alkalinity added as carbonates). Points show the relative fluorescence values of each replicate and lines show loess fits. Chl-a fluorescence (ex/em at 445/685 nm) is shown for all strains except the cyanobacteria, for which phycocyanin fluorescence (ex/em at 586/647 nm) is shown. Continued on next page.

Fig. S1 continued:

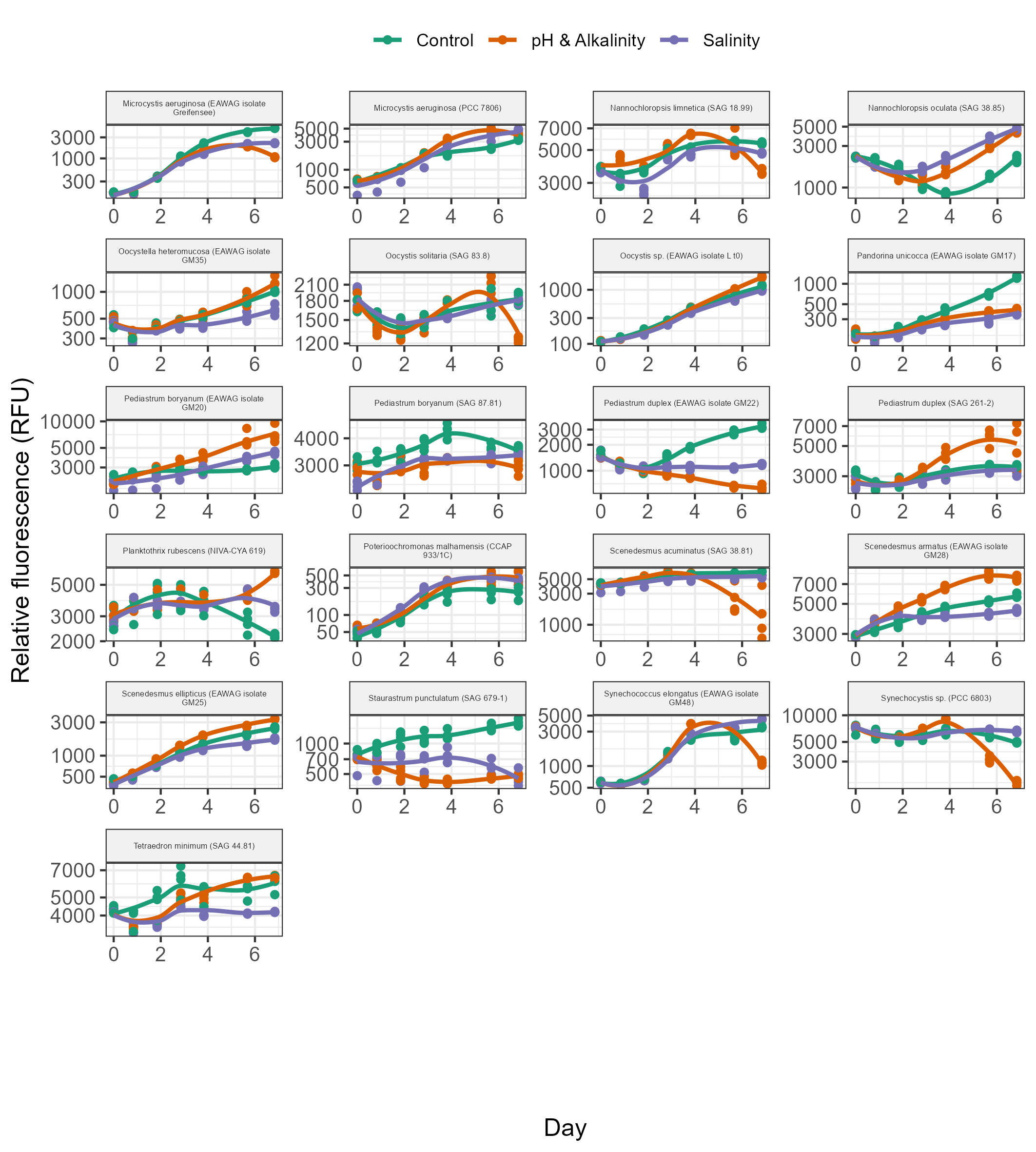

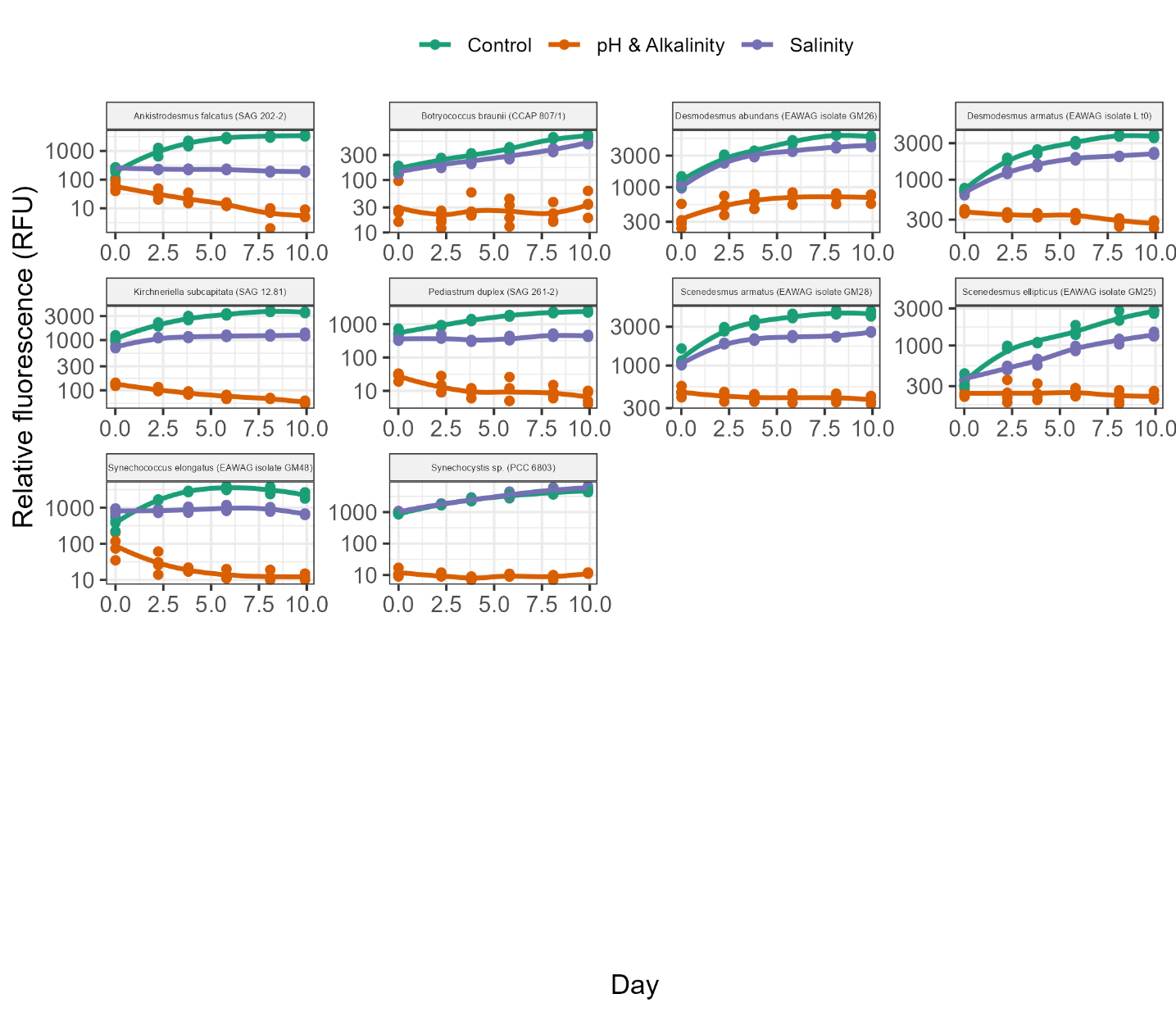

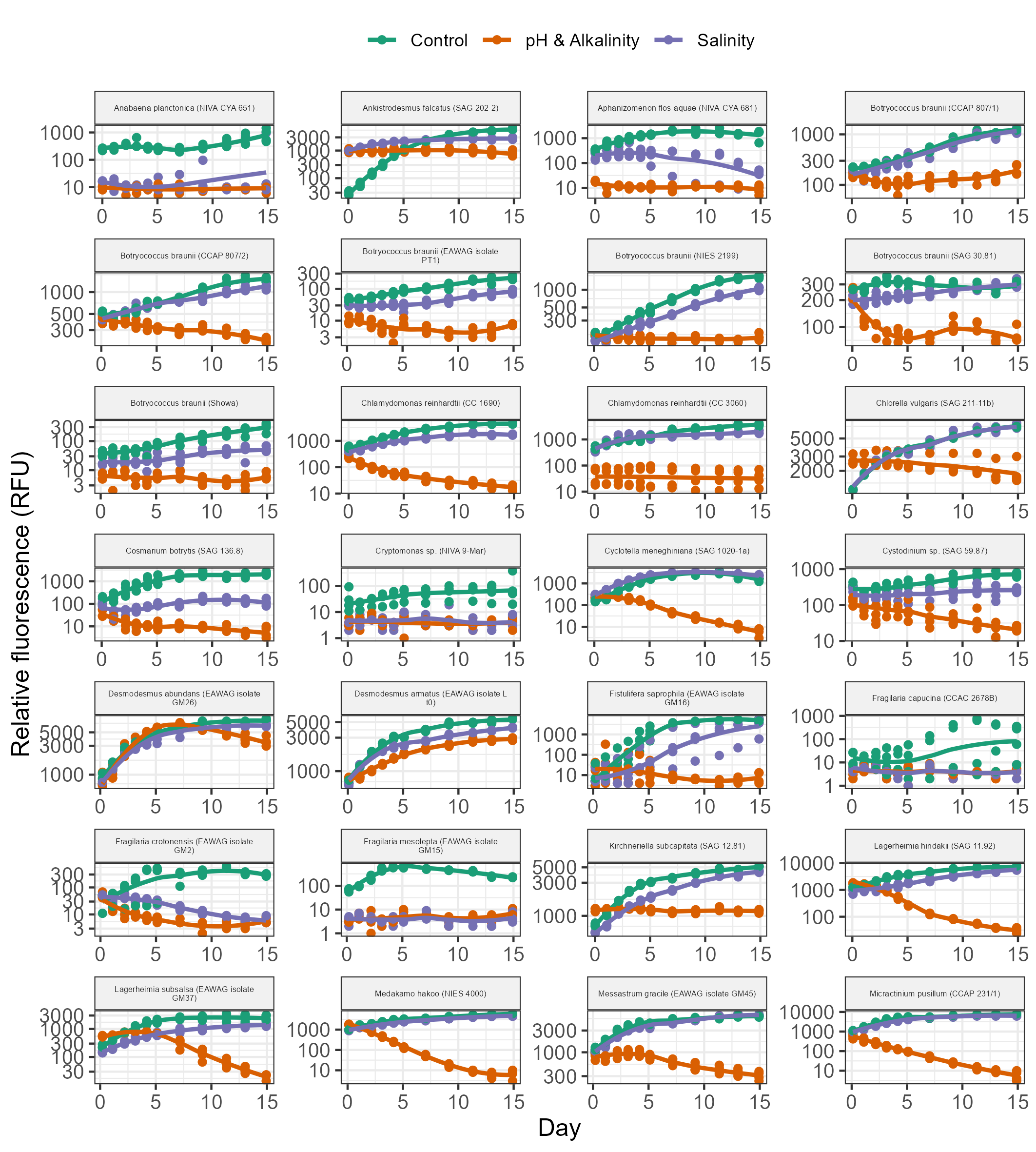

Fig. S2. Growth curves of all microalgae strains in Phase 2 (i.e., the screening in high pH and alkalinity – pH 10 and 75 mM of alkalinity added as carbonates). Points show the relative fluorescence values of each replicate and lines show loess fits. Chl-a fluorescence (ex/em at 445/685 nm) is shown for all strains except the cyanobacteria, for which phycocyanin fluorescence (ex/em at 586/647 nm) is shown. Continued on next page.

Fig. S2 continued:

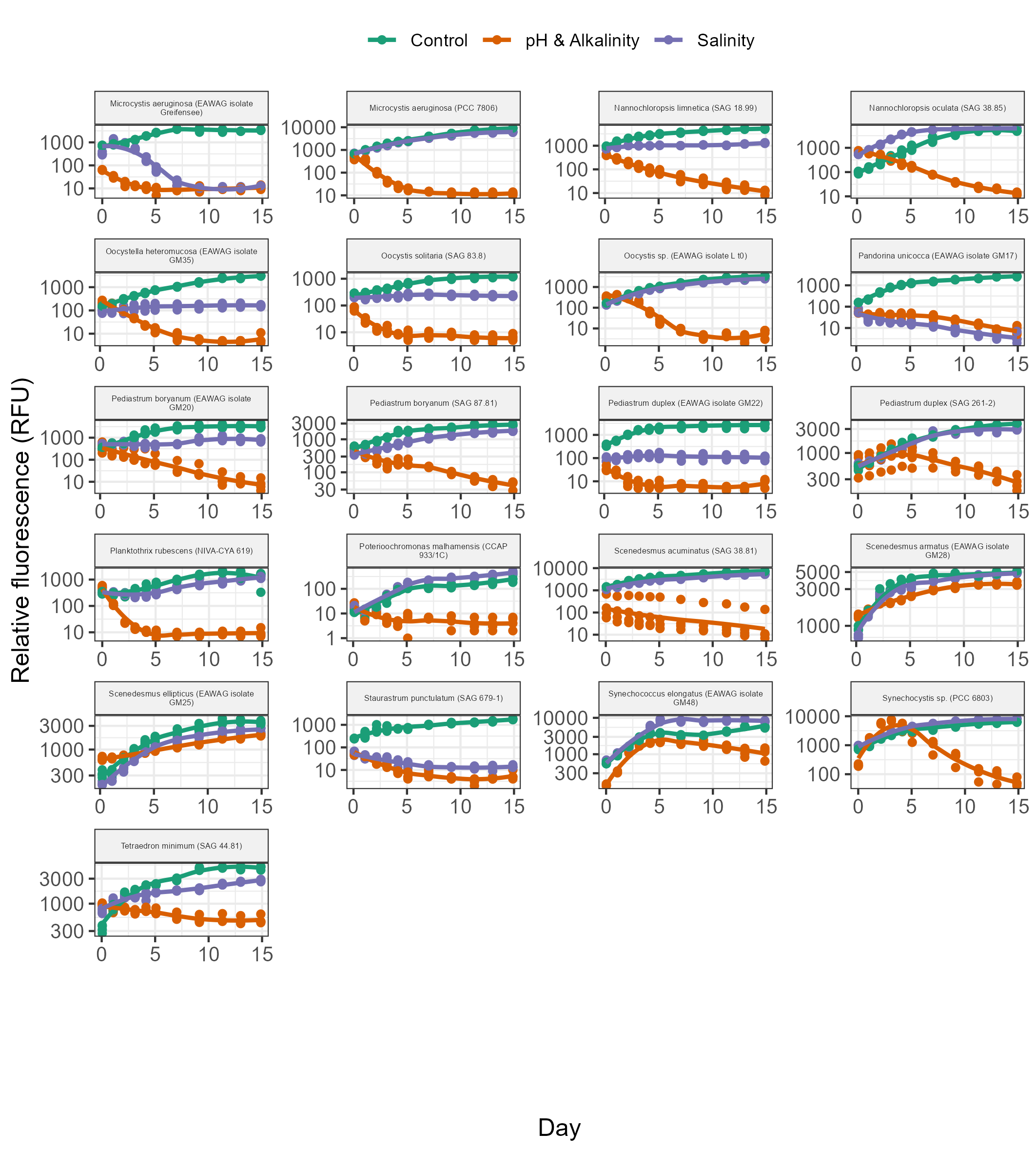

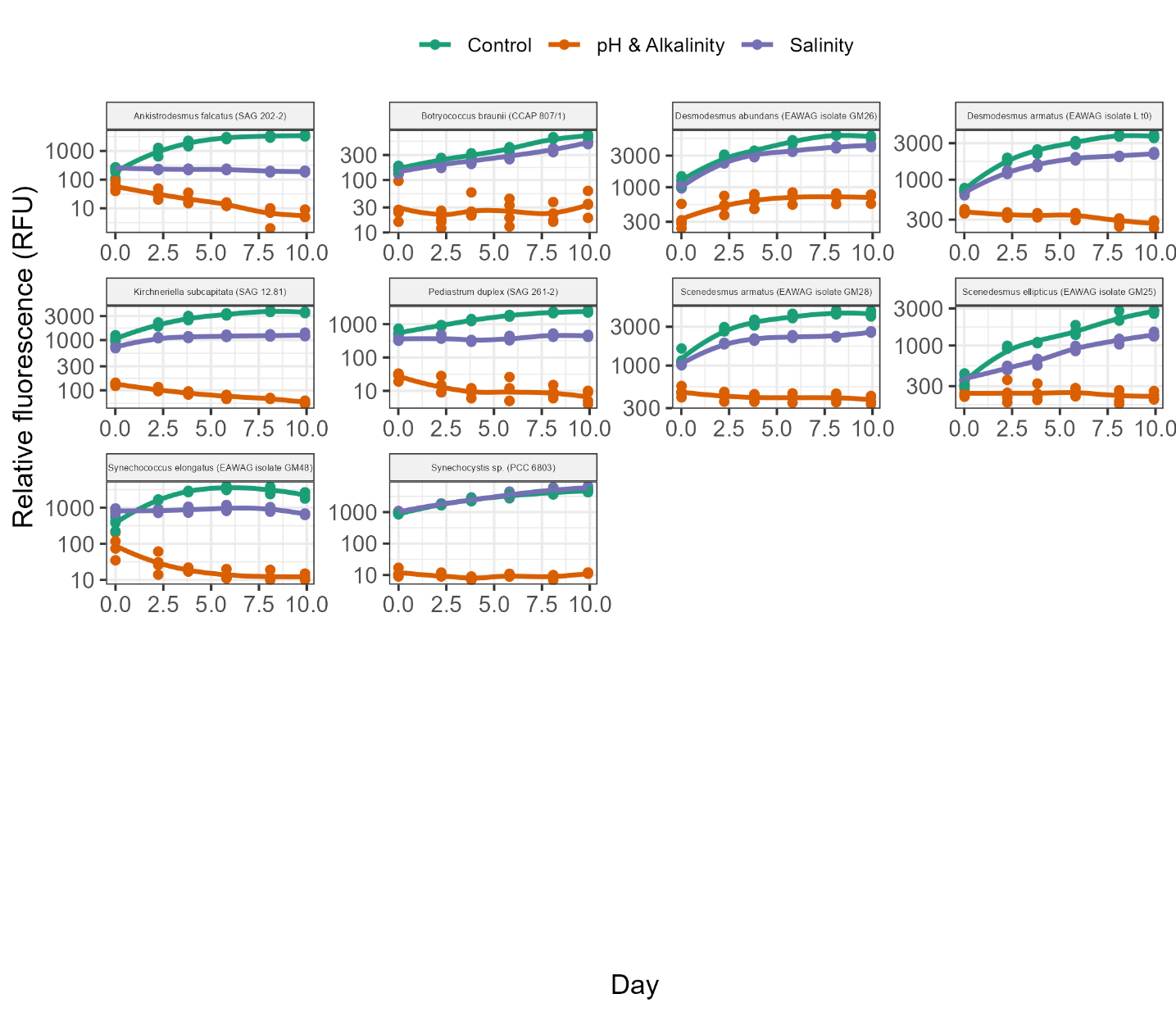

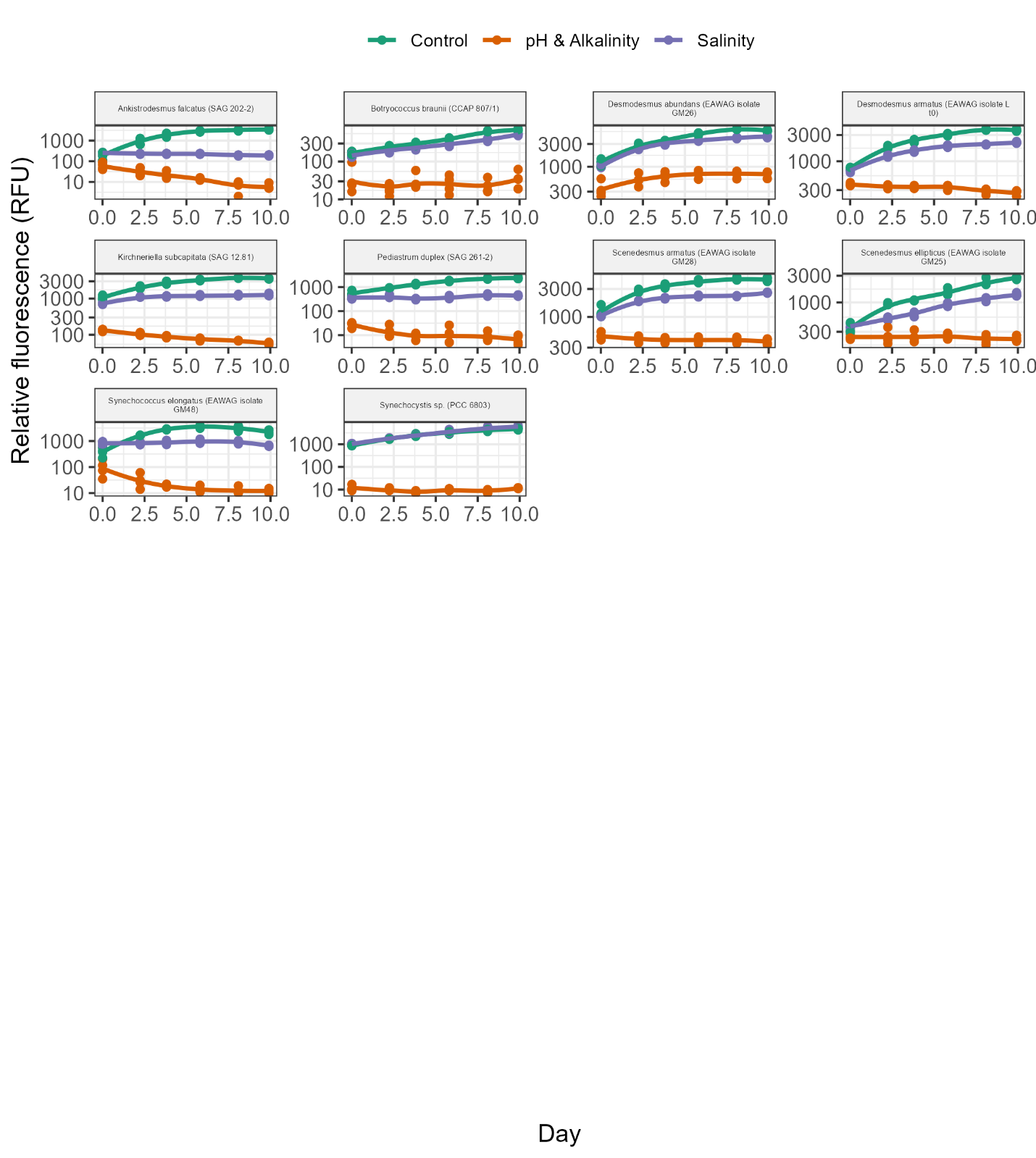

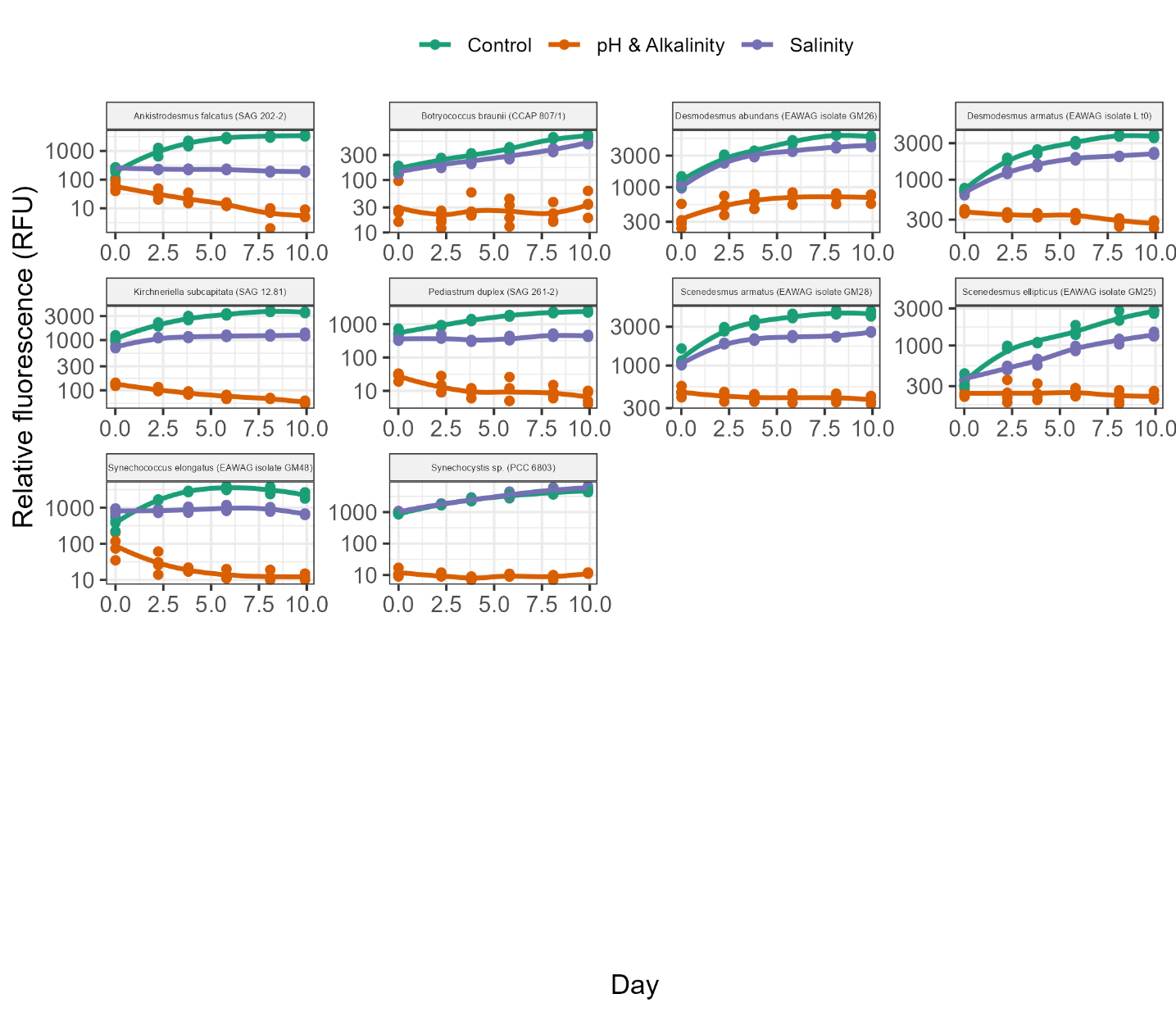

Fig. S3. Growth curves of all microalgae strains in Phase 3 (i.e., the screening in extreme pH and alkalinity – pH 10 and 150 mM of alkalinity added as carbonates). Points show the relative fluorescence values of each replicate and lines show loess fits. Chl-a fluorescence (ex/em at 445/685 nm) is shown for all strains except the cyanobacteria, for which phycocyanin fluorescence (ex/em at 586/647 nm) is shown.

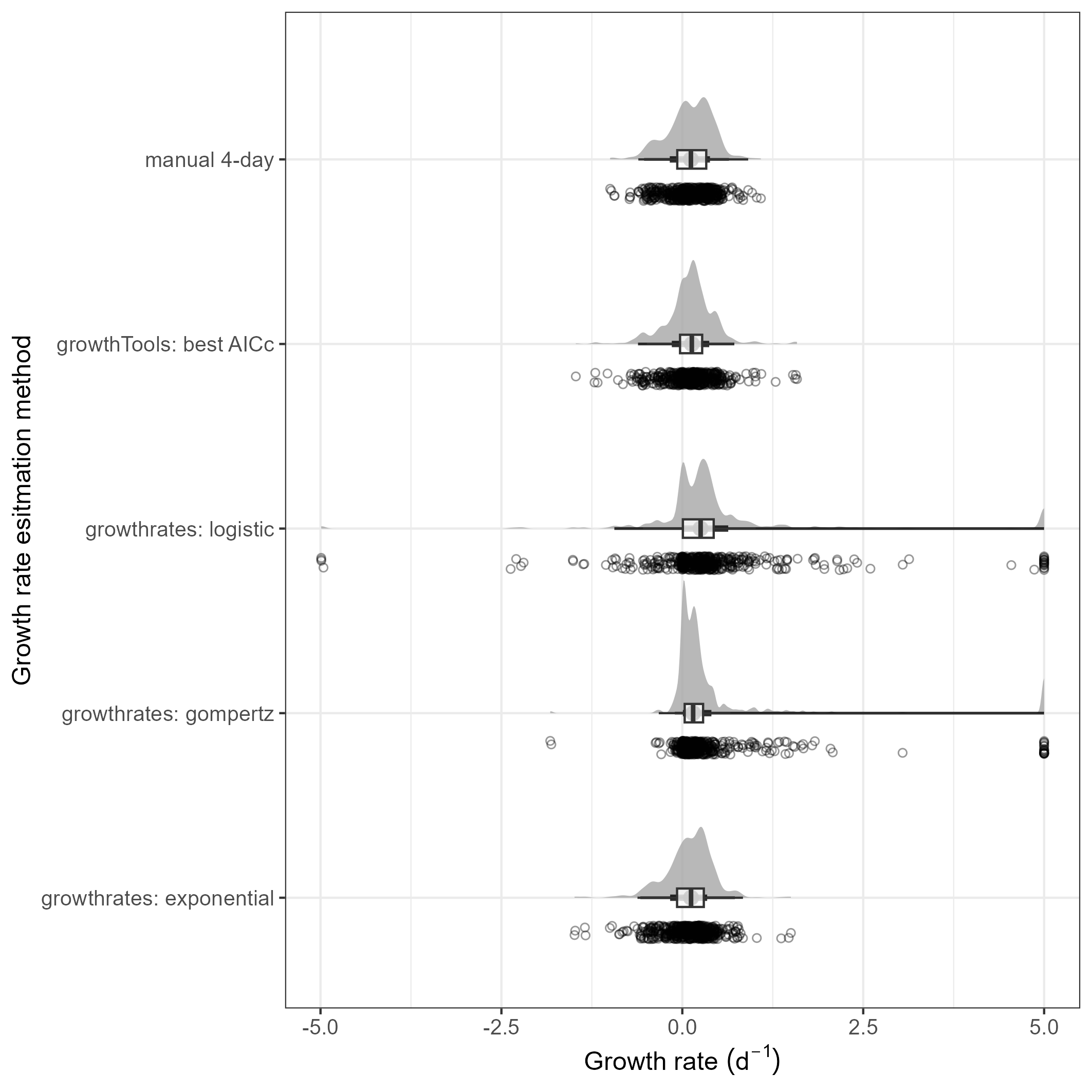

Fig S4. Distribution of growth rate estimates in Phase 2 using five different methods. Points show estimates for each experimental unit, density plots show smoothed distributions, and boxplots show default statistics (median, 25^th^ to 75^th^ percentile, and extreme data points within 1.5x the interquartile range). Logistic and Gompertz models often provided unrealistically high estimates which hit the maximum value allowed (µmax of 5), whereas the manual approach, growthTools, and growthrates (exponential) models all provided similar results.

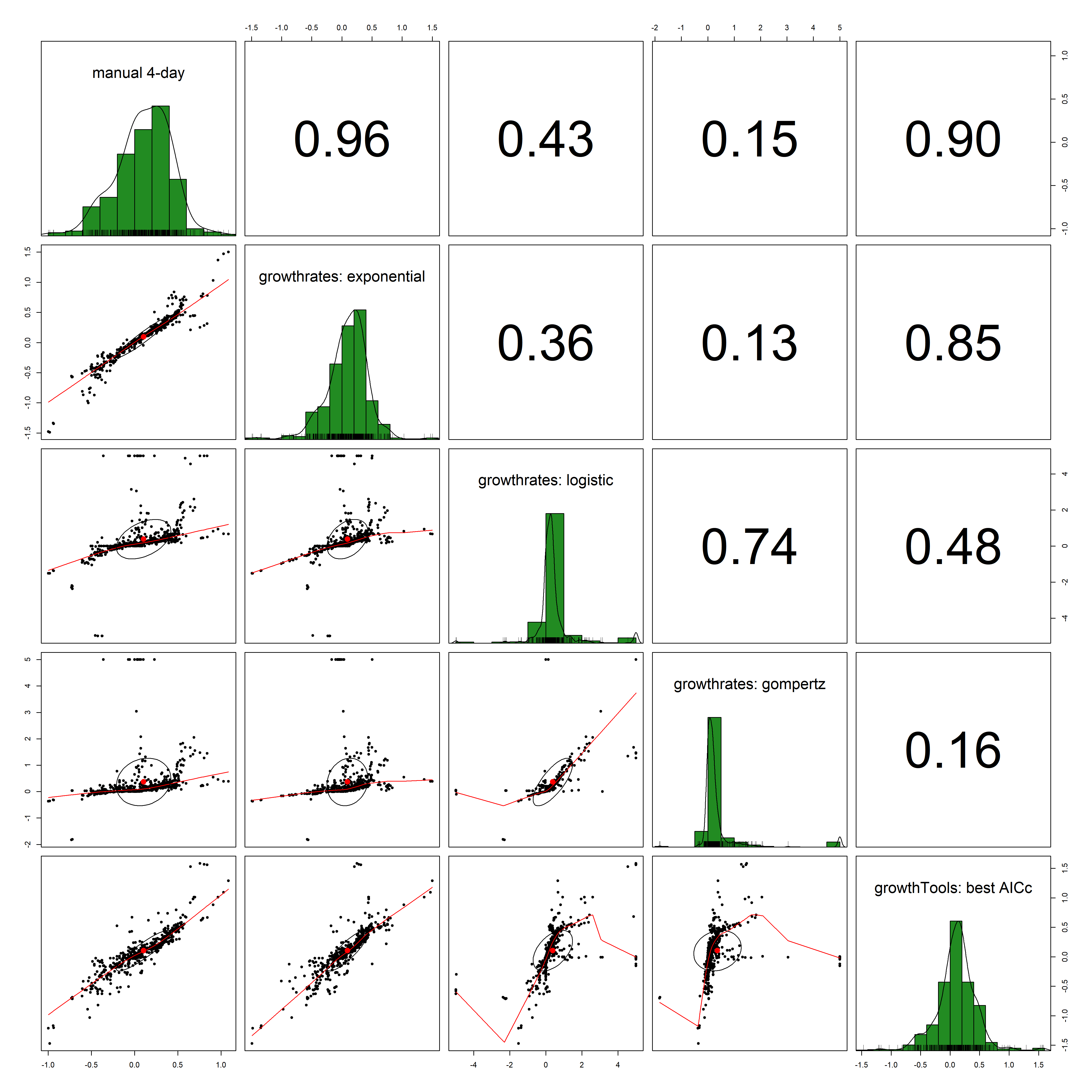

Fig S5. Correlation plots showing similarities among growth rate estimation methods applied for Phase 2. Diagonals show histograms of growth rates, numbers in upper right indicate Pearson correlation coefficients, and lower left figures plot growth rate estimate methods against each other for all 588 experimental units, with red lines showing loess fits to the data.

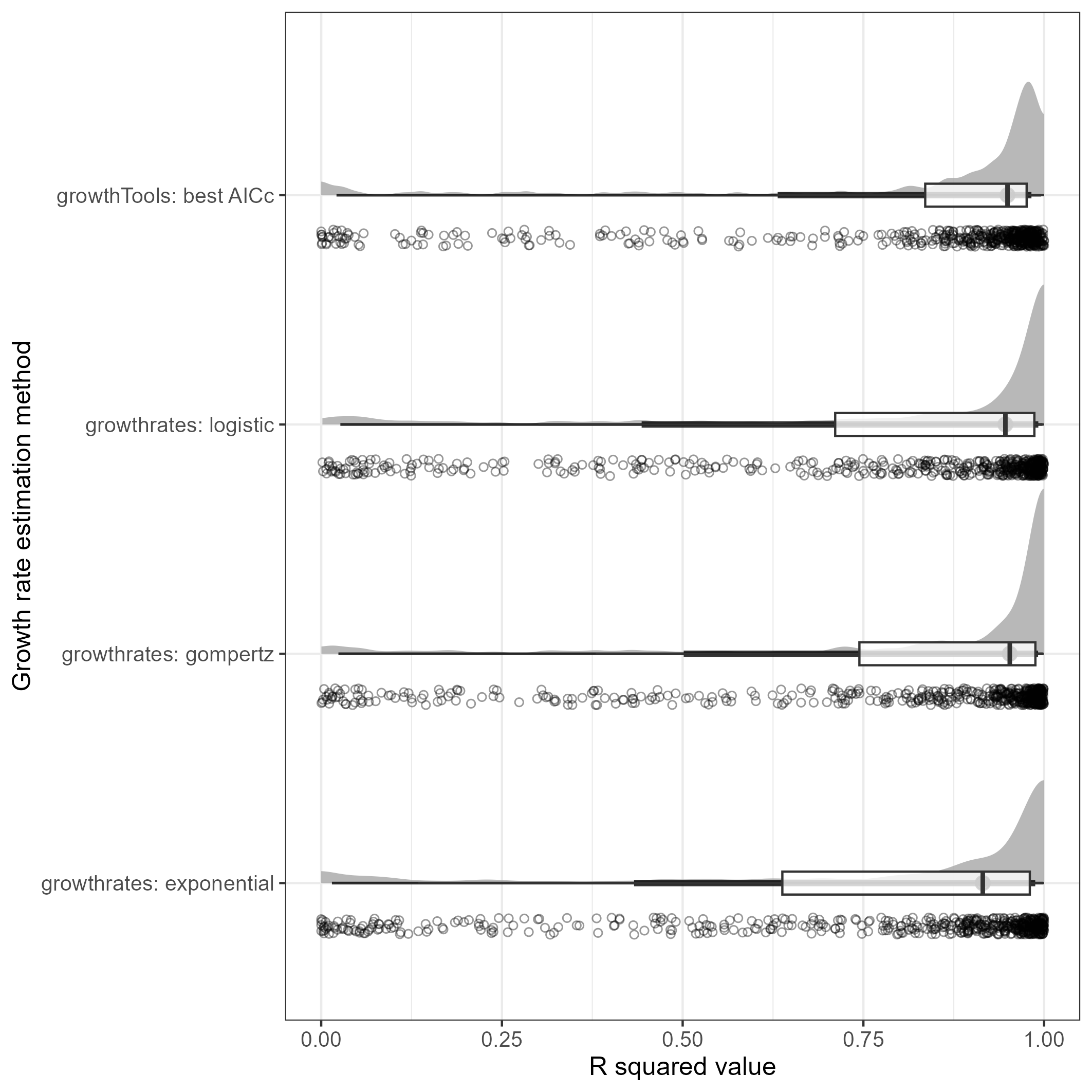

Fig S6. Distribution of R^2^ values for the four different growth models assessed in Phase 2. Points show estimates for each experimental unit, density plots show smoothed distributions, and boxplots show default statistics (median, 25^th^ to 75^th^ percentile, and extreme data points within 1.5x the interquartile range).

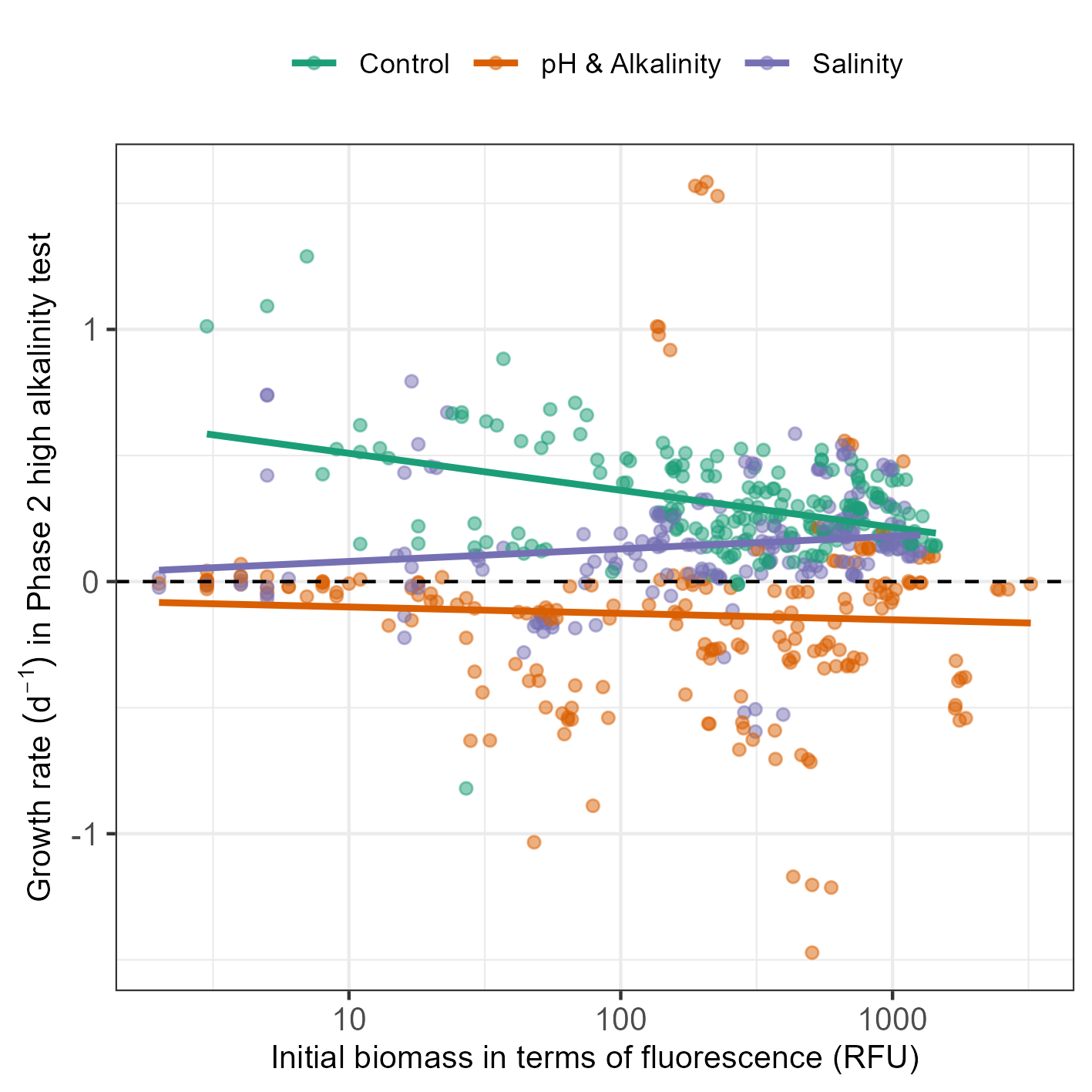

Fig. S7. Effects of the initial inoculation density (in terms of chlorophyll fluorescence) on growth rates in Phase 2. An additive linear model shows no overall effect of initial density on growth rate (p = 0.12); however, a model including an interaction term between the media treatment and initial density shows that there is a significant interactive effect; i.e., initial density effects depend on the medium type (F_5,582_ = 48.7, p-value for interaction terms < 0.007). When performing linear regressions independently for each medium type, regression parameters are as follows: Control: *µmax* = -0.064 × log_initial_density + 0.66 (R^2^ = 0.16); pH & alkalinity: *µmax* = -0.011 × log_initial_density – 0.075 (R^2^ = 0.002); Salinity: *µmax* = 0.022 × log_initial_density + 0.030 (R^2^ = 0.026). In summary, this analysis of the effect of initial densities shows no effect for the pH & alkalinity treatment, a small positive effect for the salinity treatment, and a negative effect for the control treatment. Therefore, growth rates may be underestimated for controls when transfers resulted in relatively initial high densities, but there is no evidence to suggest that growth in either the pH & alkalinity or salinity treatments was inhibited due to relatively high initial densities.
